# AmpliPhy improves gene trees by adding homologous sequences without affecting alignments

**DOI:** 10.64898/2026.01.26.701724

**Authors:** Dongwook Kim, Manuel Gil, Kazutaka Katoh, Christophe Dessimoz

## Abstract

In phylogenomics, gene tree reconstruction depends on multiple sequence alignment (MSA) and tree inference, and ongoing work continues to improve inference quality. Denser taxon sampling has been associated with improved gene tree inference, suggesting that adding homologs could be a practical route to higher accuracy as sequence databases continue to expand. However, adding sequences can influence multiple steps of typical inference pipelines, and little is known on its specific effect on the multiple sequence alignment, tree reconstruction, and rooting steps. We performed a large-scale empirical and simulated benchmarks to quantify how homolog enrichment affects alignment and phylogenetic inference. Using an enrichment–impoverishment design and a measure of tree accuracy based on taxonomic congruence, we found that enrichment consistently improves tree inference quality, while effects on alignment quality are marginal. We show that this improvement is associated with, but not restricted to accurate root placement on enriched trees when sensitive homolog search is accompanied. Notably, much of the benefit can be retained with relatively compact alignments produced by sequence addition. Building on these observations, we provide a tool, AmpliPhy, which efficiently improves phylogenetic reconstruction of protein families through homolog enrichment. The AmpliPhy open-source pipeline software is available at https://github.com/DessimozLab/ampliphy.

## Introduction

The standard phylogenomics workflow for a gene family consists of multiple sequence alignment (MSA) followed by phylogenetic tree inference (Kapli et al., 2020). Phylogenetic inference is sensitive to MSA quality and to the performance of the alignment and tree inference methods (Lozano-Fernandez, 2022; Ogden and Rosenberg, 2006), and numerous efforts have evaluated and improved gene tree reconstruction quality in this context (Dessimoz and Gil, 2010; Iantorno et al., 2014; Landan and Graur, 2007; Liu et al., 2009; Zhou et al., 2018)

Improvements in tree inference quality can be achieved by enriching an MSA with additional homologous sequences, consistent with decades of work assessing the effects of dense taxon sampling (Nabhan and Sarkar, 2012). With the continued expansion of public sequencing archives (Chikhi et al., 2025; Sayers et al., 2024), it has become increasingly feasible to retrieve large numbers of homologs per gene family and revisit enrichment as a practical route to improving phylogenetic inference. The general consensus is that increased taxon sampling benefits phylogenetic inference, reducing phylogenetic error (Dong et al., 2022; Zwickl and Hillis, 2002) and alleviating artifacts such as long-branch attraction (Ontano et al., 2021). Yet, most existing studies assess the effect of added sequences primarily at the tree inference stage, while the impact of taxon sampling on alignment quality has largely been overlooked. Moreover, taxon sampling can also affect root placement, for example through outgroup choice and the mitigation of long-branch artifacts (Graham et al., 2002; Stefanović et al., 2004). Consequently, we still lack a systematic and scalable method to decompose and evaluate how added sequences influence the alignment, tree inference, and rooting steps.

Traditionally, such evaluations have been limited by scalability and by the difficulty of applying consistent methodology at large scale. Enriching taxa can quickly inflate alignment size, increasing both alignment handling and likelihood-based tree search workloads (Sanderson and Driskell, 2003). Even when inference is feasible, defining scalable, quantitative criteria for comparing phylogenetic outcomes across many gene families remains non-trivial (Lanfear and Hahn, 2024; Pease et al., 2018; Zhou et al., 2020). However, recent improvements in the scalability of sequence alignment and maximum-likelihood inference make large-scale benchmarks increasingly feasible (Deorowicz et al., 2016; Edgar, 2022; Katoh and Standley, 2013; Kozlov et al., 2019; Minh et al., 2020). Among these, MAFFT provides a practical route to enrichment by inserting newly found sequences into an existing alignment without recomputing the full MSA from scratch (Katoh and Frith, 2012).

Recent methodological advances in phylogenetic tests are also relevant, notably the enriched and impoverished species discordance test (Gil, 2010). This framework separates the effect of adding taxa on alignment quality from its effect on tree inference. The phylogenetic tree inferred from the *enriched* alignment—an alignment of the combined set of the original sequences and additional homologs—is jointly influenced by (i) changes in the sequence alignment and (ii) the contribution of additional taxa to phylogenetic inference. In contrast, the *impoverished* alignment, in which the additional taxa are removed from the enriched MSA, yields a phylogenetic tree that reflects only the effect of enrichment on alignment quality. Comparing trees inferred from original, enriched and impoverished MSAs thus isolates the contribution of additional taxa to tree inference and sequence alignment (see Fig. 1).

**Fig 1.**
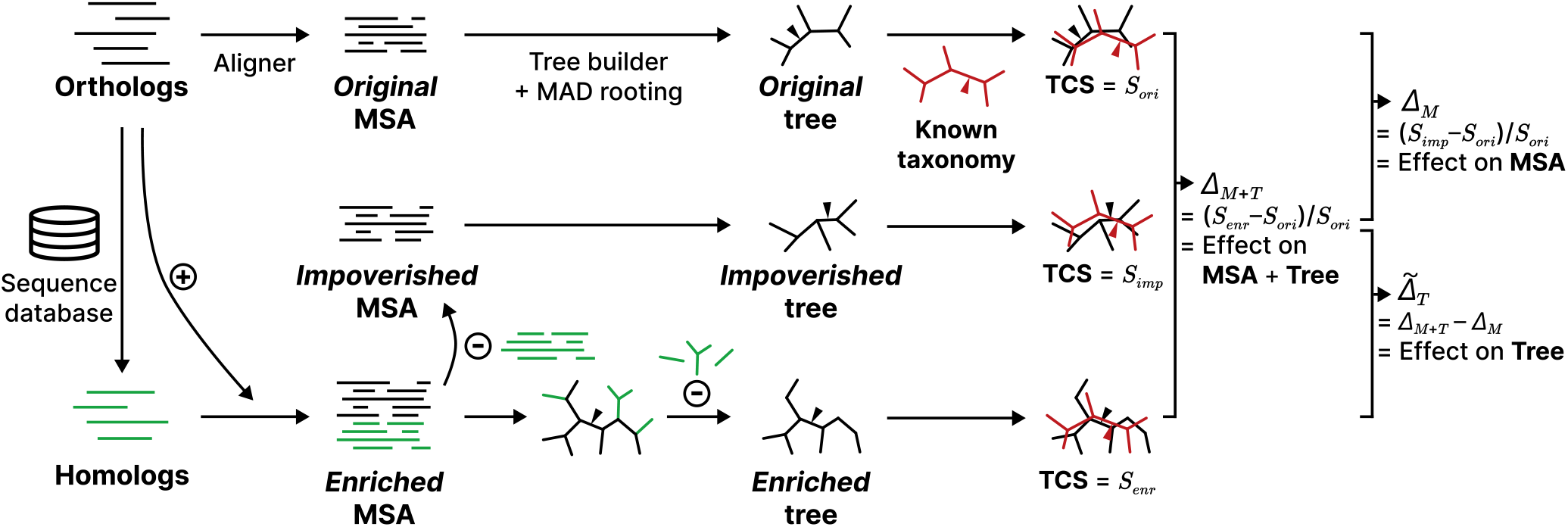
Enriched–impoverished testing of MSAs combined with taxonomic congruence decomposes the effects of homolog enrichment on phylogenetic inference. For each orthologous gene family, we constructed three MSAs: *original*, computed by applying sequence aligners directly to the input sequences; *enriched*, computed by aligning the combined set of orthologs and homologs identified by database search; and *impoverished*, obtained by removing the added homologs from the enriched MSA. We then used TCS to quantify congruence of the resulting trees against the known taxonomy. The normalized difference in congruence between the original and enriched trees, denoted Δ_*M*+*T*_, captures the joint impact of sequence addition on alignment and tree inference. Δ_*M*_, the normalized difference between the original and impoverished trees, reflects the effect on alignment quality alone. The effect on tree inference can then be estimated by subtraction, 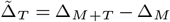 .

Moi and colleagues (Moi et al., 2025) recently revisited taxonomic congruence as a measure of phylogenetic inference quality for large-scale structural phylogenetics. The taxonomic congruence score (TCS; Tan et al., 2015) measures how well a gene tree agrees with an external species taxonomy and provides a quantitative criterion for evaluating trees via agreement with an established species phylogeny (Dessimoz and Gil, 2010).

Building on these advances, we conducted a large-scale empirical benchmark by combining the enrichment– impoverishment framework with TCS, decomposing the effect of homolog enrichment on alignment, tree inference, and rooting (Fig. 1). We show that homolog enrichment consistently improves phylogenetic inference quality, with the improvement associated with accurate root placement on enriched trees of closely related taxa, while changes in alignment quality are marginal. We further show that much of this benefit can be retained with relatively compact alignments produced by sequence addition compared with full realignment. Finally, we present an easy-to-use pipeline that automates homolog enrichment and improves phylogenetic re-construction for gene families.

## Methods

### Benchmark species and orthologous group selection

We obtained the eukaryotic subset of the Quest for Orthologs (QfO) 2024_02 reference proteomes (Langschied et al., 2024), which comprises 51 species. One species (*Daphnia magna*) was not available in the OMA release All.Jul2024 (Altenhoff et al., 2024) and was therefore excluded. From the root hierarchical orthologous groups (root HOGs) in the OMA database (Altenhoff et al., 2019), we selected 100 random groups with fewer than 50 member sequences while representing at least 10 species. We applied the same selection procedure of HOGs to other taxon sets (Amniota, Primates, Bacteria, Actinomycetes, and larger Eukaryotic HOGs; see Supplementary Table 1 for details).

### Homolog enrichment and MSA impoverishment

We used the UniRef50 database (UniProt clustered at 50% sequence identity) from UniProt release 2025_02 (UniProt Consortium, 2025) to identify homologs for the selected HOGs. For each HOG, we ran the `easy-search` module of MMseqs2 v17.b804f (Steinegger and Söding, 2017) with different *E*-value thresholds (`-e 1e-100, 1e-10, 1e-3, 1e-1, 1e0`) to define homologous sequences from the database. We then extracted, for each homolog, the subsequence locally aligned to the original sequence and randomly sampled up to 5× the size of the original HOG sequence set. We combined these homologous subsequences with the original sequences and aligned the combined set using multiple sequence aligners (Deorowicz et al., 2016; Edgar, 2022; Katoh and Standley, 2013; Katoh and Toh, 2008; Larkin et al., 2007; Sievers et al., 2011; see Supplementary Table 2 for the full list of aligners used in this study), producing *enriched* MSAs. From each enriched alignment, we extracted the subset corresponding to the original sequences, which we defined as *impoverished* MSAs. We also computed MSAs directly from the initial HOG sequences, which we denote as *original* MSAs.

### Stratification of the homologs for enrichment

As a counterpart to the cumulative definition of homologous sequences under increasingly relaxed *E*-value thresholds, we implemented a stratified analysis of homologs to decompose the effect of homolog addition. For each Amniota HOG, we stratified the homologs identified with an *E*-value threshold of `-e 1e0` into four quartiles based on their local alignment bitscore. Each stratum was processed using the same procedure described above, including subsequence extraction, random sampling, enrichment, and impoverishment.

We next stratified the homologs based on their fractional identity in the alignment. For an aligned query sequence *Q* = *q*_1_ … *q*_*n*_ and target sequence *T* = *t*_1_ … *t*_*n*_, fractional identity of *T* given *Q* can be defined as following:

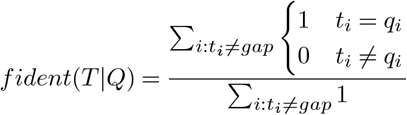

We divided the resulting identity values *x* into seven buckets: 30% ≤ *x* < 40%, 40% ≤ *x* < 50%, …, and 90% ≤ *x* ≤ 100%. Each bucket was treated as a separate stratum for sampling and enrichment.

Finally, we stratified homologs by their taxonomic relationship to Amniota. Based on the NCBI taxonomy assignment of each UniRef entry, homologs were classified as intra-clade, inter-clade, inter-phylum, inter-kingdom, or inter-domain, corresponding respectively to Amniota, non-amniote Chordata, non-chordate Metazoa, non-metazoan Eukaryota, and non-Eukaryota. Each category was treated as a separate stratum and subjected to the same analysis.

### Tree inference and calculation of the taxonomic congruence

Each HOG has a set of homologs defined at distinct *E*-value thresholds, and each HOG–homolog set yields three versions of MSAs: original, enriched, and impoverished. We ran IQ-TREE v2.4.0 (Minh et al., 2020) to infer phylogenies from these MSAs under an identical evolutionary model (`JTT+I+G4`) to enable consistent comparison of inferred topologies. We additionally used three models (`LG+I+G4`, `WAG+I+G4`, `MF`) to test the robustness of our approach across evolutionary models.

All resulting trees were rooted using minimum ancestor deviation (MAD) as implemented in MAD v2.2 (Tria et al., 2017). For trees inferred from enriched MSAs, which include additional leaves from homologs, we used GoTree v0.4.5 (Lemoine and Gascuel, 2021) to prune them to sub-trees containing only leaves corresponding to the original MSA. Additionally, we generated subtrees from enriched MSAs by switching the order of pruning and MAD rooting (see Section for details). We also inferred phylogenetic trees of Eukaryota HOGs using faster alternative tree builders while using MAFFT as a sequence aligner, including FastTree v2.1.11 with and without `--fastest` option (Price et al., 2010), and IQ-TREE with `--fast` option.

We used the taxonomic congruence score (TCS; Tan et al., 2015) to assess the resulting phylogenetic trees. The algorithm for computing TCS for a phylogenetic tree given a species tree is described in Moi et al., 2025. For Eukaryota and Amniota HOGs, we computed TCS against the reference taxonomy tree from the NCBI Taxonomy Database (Cox et al., 2025). For Bacteria and Actinomycetes HOGs, we used the taxonomy tree from Genome Taxonomy Database Release 10 (Parks et al., 2026). We normalized the scores of trees inferred from enriched and impoverished MSAs by dividing them by the score of the tree inferred from the corresponding original MSA (Δ_*M*+*T*_ and Δ_*M*_, respectively. See Fig. 1 for details).

We used one-sided paired *t*-tests to assess the statistical significance of differences in normalized TCS gains. For comparisons between enriched and impoverished TCS values (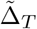, Fig. 2c,f), we tested the null hypothesis *µ* ≤ 0. For comparisons between pre- and post-pruned trees (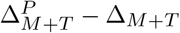, Fig. 3c,d), we tested the null hypothesis *µ* ≥ 0. Resulting *p*-values were adjusted using the Benjamini–Hochberg procedure to control the false discovery rate (Benjamini and Hochberg, 1995).

**Fig 2.**
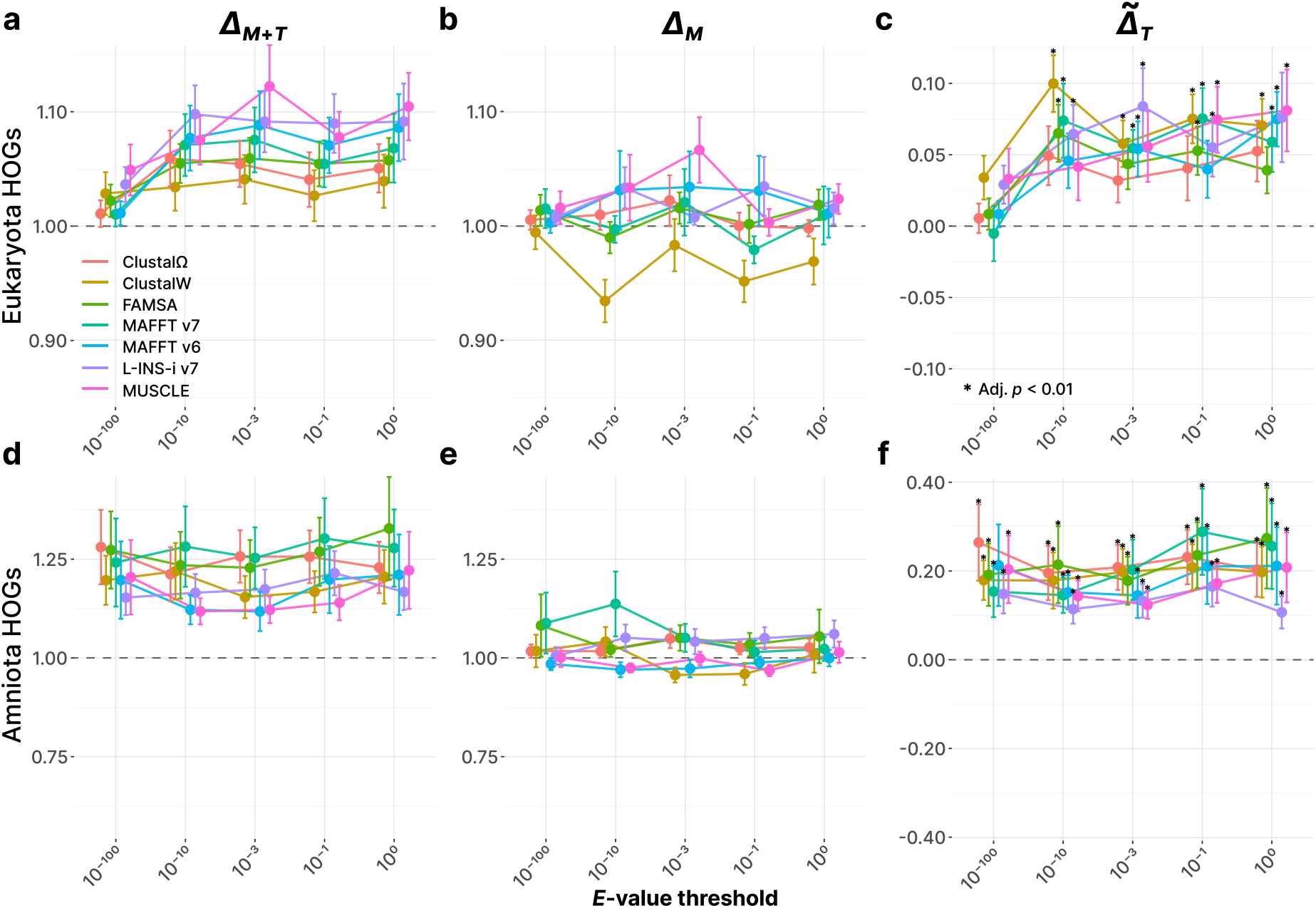
Homolog enrichment of multiple sequence alignments improves tree quality, but has marginal effects on alignment quality. Dashed lines indicate the threshold between positive (above, more congruent) and negative (below, less congruent) estimated impact. Data points are colored by the MSA program used to compute the alignment. **a**. Mean standard error of the normalized TCS differences for enriched trees (Δ_*M*+*T*_ ; impact of sequence addition on both alignment and tree building) from 100 Eukaryota HOGs. **b**. Mean standard error of the normalized TCS differences for impoverished trees (Δ ; impact on alignment quality). **c**. Difference between Δ_*M*+*T*_ and Δ_*M*_ (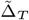 ; impact on tree quality). Statistically significant points (BH-adjusted *p <* 0.01) are annotated with asterisks. **d–f**. Corresponding taxonomic congruence statistics from a relatively closely related set of species (100 Amniota HOGs).

**Fig 3.**
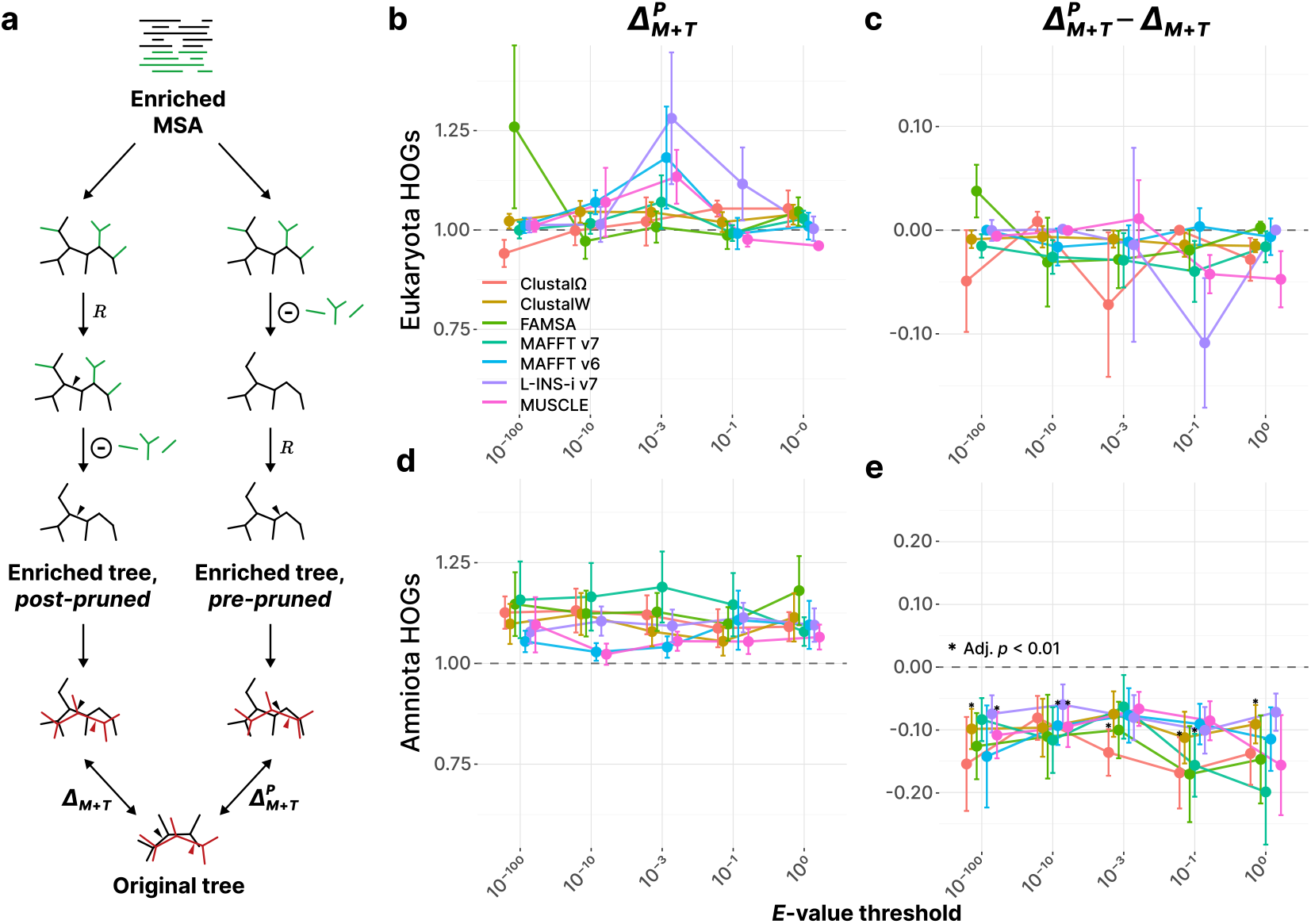
Sensitive homolog search improve tree inference quality by enabling more precise root placement. a. Schematic illustration of the post- and pre-pruning procedures for enriched trees. The process denoted by *R* indicates rooting by minimum ancestor deviation. We computed the overall gain in taxonomic congruence for trees built with pre-pruning 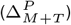 . **b**. Normalized TCS gain for pre-pruned trees inferred from enriched MSAs 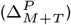 of 100 Eukaryota HOGs. **c**. Loss of congruence due to pre-pruning 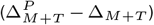. Asterisks indicate statistically significant points (BH-adjusted *p <* 0.01). **d–f**. Corresponding taxonomic congruence statistics for a more closely related taxon set (100 Amniota HOGs).

### Simulation-based benchmark of the enrichment– impoverishment design

We performed a simulation to further benchmark the plausibility of our enrichment–impoverishment design. Using ALF v0.99 (Dalquen et al., 2012), we simulated the evolution of 500 species, each with a genome comprising genes from 100 orthologous families. We then randomly selected 50 species and obtained the corresponding 100 orthologous gene families. For each family, homologous sequences for enrichment were retrieved by searching the sequences from these 50 species against those from the remaining 450 species. Using these sequences, we generated original, enriched, and impoverished alignments as described above.

We calculated the Robinson–Foulds (RF) similarity of the original (*T*_*o*_), enriched (*T*_*e*_), and impoverished (*T*_*i*_) trees to the ground-truth evolutionary tree (*T*_*g*_) for the sampled 50 species (Robinson and Foulds, 1981). For two trees *T*_1_ and *T*_2_ with shared number of leaves *n* and RF distance *d*, RF similarity was calculated as:

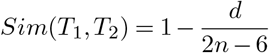

The combined impact of enrichment on the MSA and tree inference steps (*RF*_*M*+*T*_ ) was measured as *Sim*(*T*_*e*_, *T*_*g*_) −*Sim*(*T*_*o*_, *T*_*g*_). Similarly, the impact on the MSA step alone (*RF*_*M*_ ) was measured as *Sim*(*T*_*i*_, *T*_*g*_) −*Sim*(*T*_*o*_, *T*_*g*_). The estimated impact on the tree inference step (*RF*_*T*_ ) was then defined as *RF*_*M*+*T*_ − *RF*_*M*_ .

### Adding homologs to a pre-computed MSA

Profile alignment methods, such as those implemented in ClustalW, allow new sequences to be aligned to an existing multiple sequence alignment, thereby defining homologous regions relative to a pre-aligned sequence family (Thompson et al., 1994). MAFFT implements this feature at scale, by adding sequences while preserving the column structure of the input MSA via the `--add` and `--addfragments` options, accompanied with `--keeplength` option (Katoh and Frith, 2012). We used these options to add the selected homologs to the original MSA, either with higher sensitivity and *O*(*N* ^2^) time complexity (`--add`) or with linear scalability (`--addfragments`). We refer to MSAs constructed in this way as *amplified*, to distinguish them from enriched MSAs. We performed phylogenetic inference, rooting, pruning, and TCS calculation for amplified MSAs as described above.

## Results

### Homolog enrichment of MSA improves phylogenetic inference by impacting tree quality

We computed taxonomic congruence scores against the NCBI taxonomy tree (Cox et al., 2025) for phylogenetic trees inferred from 100 eukaryotic hierarchical orthologous groups (HOGs; Jothi et al., 2006) from the OMA database (Altenhoff et al., 2024). Fig. 2a and b respectively show the effect of sequence addition both on MSA and tree (Δ_*M*+*T*_, normalized TCS difference between enriched and original MSAs) and the effect on MSA only (Δ_*T*_, between impoverished and original MSAs). We gradually relaxed the *E*-value thresholds used to define homologs, from very stringent (10^−100^) to very permissive (10^0^) values.

By subtracting these scores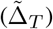, we observed a positive impact of homolog enrichment on phylogenetic tree inference step, regardless of the sequence aligner used to build the alignments (Fig. 2c). This effect was even more pronounced from HOGs of relatively closely related species (Amniota, Fig. 2d–f), possibly due to the database homolog searches becoming more sensitive when the query taxa are more closely related (Pearson, 2013). Notably, both in Eukaryota and Amniota datasets, the impact of sequence addition on alignments (Δ_*M*_ ) was marginal.

We observed the consistent results from the analysis using a more closely related set of taxa (order Primates, Supplementary Fig. 1), as well as with a larger Eukaryotic HOGs with up to 200 members (Supplementary Fig. 2). The results from Amniota HOGs were also consistent across different choices of evolutionary models, such as `LG+I+G4`, `WAG+I+G4`, or `MF` (Supplementary Fig. 3).

Additionally, we assessed the effect of enrichment on bacterial HOG trees using the current GTDB taxonomy (Parks et al., 2026). We observed improved phylogenetic tree inference for enriched Actinomycetes HOGs (Supplementary Fig. 4a–c); however, the difference was marginal for HOGs spanning the entire domain Bacteria (Supplementary Fig. 4d– f). This may be due to the broader taxonomic scope, which can reduce homolog search sensitivity, and to inconsistencies in bacterial HOGs caused by frequent lateral gene transfer (Sarton-Lohéac et al., 2025). Here, we used the GTDB taxonomy instead of the NCBI taxonomy tree because TCS showed lower resolution when calculated using the NCBI tree (Supplementary Fig. 5), indicating that a robust taxonomic lineage definition improves the accuracy of TCS-based evaluation.

To decompose and quantify the effect of sequence addition, we stratified homologs by alignment score, sequence identity, and taxonomic distance. Stratification by alignment score or sequence identity revealed no clear differences in improvement across strata (Supplementary Fig. 6a–b). In contrast, stratification by taxonomic relationship showed that the addition of taxonomically close sequences contributed most to the improvement, whereas the effect diminished as the added sequences became more taxonomically distant (Supplementary Fig. 6c). These results suggest that homolog addition may improve gene tree inference primarily by enriching local taxonomic sampling, rather than through a specific dependence on sequence-space distance.

### Improvement of tree quality by homolog enrichment is consistent under simulated evolution

We further confirmed our observations from the empirical dataset in a simulated setting. We compared the topologies of the original, enriched, and impoverished gene trees against the ground-truth evolutionary history of each gene using Robinson–Foulds distances. Consistent with the empirical dataset, homolog enrichment had a prominent impact on tree inference (*RF*_*T*_ ), whereas its impact on the MSA step (*RF*_*M*_ ) remained marginal (Supplementary Fig. 7).

In addition, we generated taxonomic lineages from the ground-truth gene trees by enumerating their internal nodes, which were then used to calculate TCS values. The resulting patterns were consistent with the observations based on Robinson–Foulds similarities (Supplementary Fig. 8). Taken together, these results further support the hypothesis that homolog enrichment improves tree inference quality, and also reinforce the validity of TCS as an indicator for quantifying tree quality.

### Estimated tree quality measure favors more exhaustive tree search

We assessed whether our estimated measure of tree inference quality 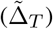 behaves as expected by comparing tree building methods while holding the aligner fixed to MAFFT. Compared with faster heuristic alternatives such as FastTree 2 (Price et al., 2010) or IQ-TREE 2 with the `--fast` option, IQ-TREE 2 under default settings inferred trees with higher taxonomic congruence (Supplementary Fig. 9). This pattern is consistent with previous observations showing that more exhaustive strategies, rather than faster shortcuts, achieve trees with higher likelihood and topological accuracy (Zhou et al., 2018). These results therefore support the validity of our benchmark using 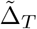 as a measure of tree inference quality.

### Sensitive homolog enrichment allows for precise root placement on the tree

We hypothesized that the gain in inference quality could be driven by more precise root placement, leveraging the additional information provided by the enriched taxa. To test this, we modified the processing of the enriched tree by switching the order of rooting and pruning (Fig. 3a). In our original workflow, we processed trees inferred from enriched MSAs by rooting first and then pruning leaves that were not present in the original MSA (*post-pruned*). By reversing the order, i.e. pruning first and rooting afterwards, we prevented the additional taxa from contributing to the rooting step (*pre-pruned*).

We then compared the congruence gain of pre-pruned 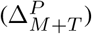 and post-pruned (Δ_*M*+*T*_ ) phylogenetic trees (Fig. 3b–e). To quantify the effect of pre-pruning, we computed the loss of congruence by pre-pruning the tree, which can be calculated as 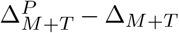 . The effect of pre-pruning was marginal for Eukaryota gene families (Fig. 3c), but we observed a notable decrease in inference quality for Amniota HOGs (Fig. 3e). Together, these results suggest that the additional taxonomic and phylogenetic information provided by sensitive homolog searches improves root placement, yielding trees with higher congruence to the established taxonomy.

### Tree improvement persists after removing the dependency on the root placement

Complementing the previous analysis of root placement, we removed the dependence on a specific rooting by selecting the best-scoring root placement across all branches (Supplementary Fig. 10a). We found that the improvement in tree quality persisted even after accounting for optimal root placement in Amniota HOGs (Supplementary Fig. 10b–d). This pattern demonstrates that accurate root placement is not the sole driver of the taxonomic improvement resulting from homolog enrichment. Together with our taxonomic stratification analysis of the added homologs, this observation suggests that such improvement may arise from an interplay between denser sampling of local taxonomy and more accurate root placement.

### Addition of homologs to a pre-computed MSA enables compact enrichment

Enriched MSAs are constructed by combining the original sequences of a gene family with their homologs, which can produce sparse and very large alignments, especially under permissive search thresholds. The computational cost of maximum-likelihood phylogenetic inference from these MSAs can increase steeply with alignment length, compounded by the number of taxa (Felsenstein, 1981), making this approach inefficient in practice. To mitigate this issue, we tested whether we could retain the signal introduced by homologs while controlling the size of the input MSA.

We used a feature of the MAFFT software suite that adds sequences while keeping the column structure of the original MSA intact, implemented via the `--add` and `--addfragments` options (Katoh and Frith, 2012). We refer to MSAs generated with this feature as *amplified* MSAs. Comparisons of normalized TCS gains (Δ_*M*+*T*_ ) between enriched and amplified MSAs showed comparable improvements in phylogenetic inference, regardless of the methodology (Fig. 4). This suggests that we can preserve linear scalability for both sequence alignment and phylogenetic inference, while retaining the improvement in inference quality provided by the additional sequences.

**Fig 4.**
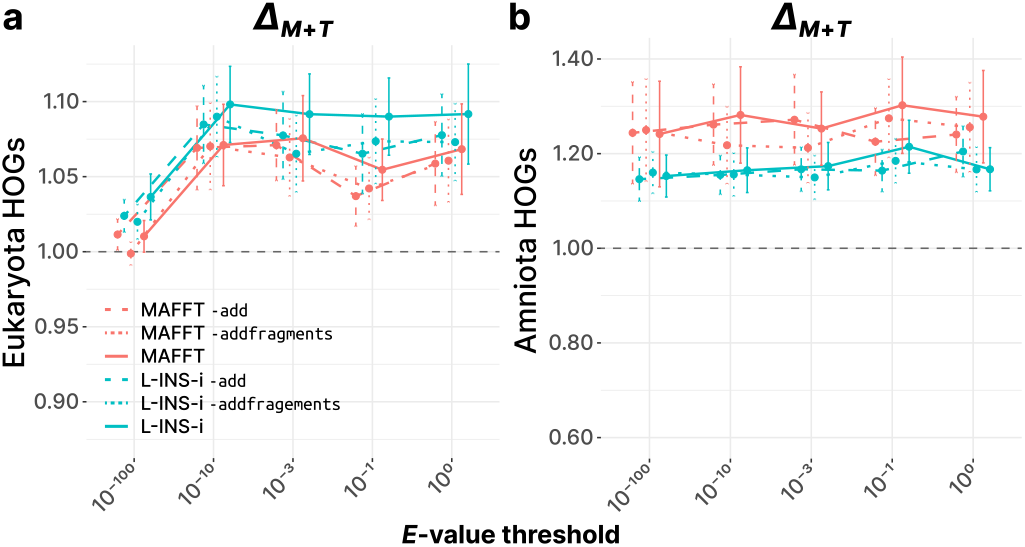
Sequence addition to a pre-computed MSA improves taxonomic congruence to a level comparable to full enrichment. a. Comparison of normalized TCS gains (Δ_*M*+*T*_ ) for trees from 100 Eukaryota HOGs inferred from enriched MSAs (solid lines), amplified MSAs generated with MAFFT `--add` (dashed lines), and amplified MSAs generated with MAFFT `--addfragments` (dotted lines). **b**. Corresponding comparison for 100 Amniota HOG trees.

### Pipeline development

We developed a pipeline named *AmpliPhy*, a scalable and fully automated solution designed to obtain improved phylogenetic trees via database homology search. AmpliPhy takes as input a set of sequences, where each set comprises homologous sequences from a gene family. This modular pipeline streamlines MMseqs2-based homolog search (Steinegger and Söding, 2017) against a reference database, alignment and amplified MSA construction with MAFFT (Katoh and Frith, 2012; Katoh and Standley, 2013), and subsequent phylogenetic inference using IQ-TREE 2 (Minh et al., 2020).

The threshold for homolog definition is fully customizable, both through alignment stringency (e.g., *E*-value, sequence identity, alignment coverage) and through an explicit limit on the relative or absolute number of sequences added per family. The pipeline reports the number of sequences added to each MSA and, when the default UniRef50 database is used as the target database, summarizes their taxonomic profile. If taxonomic metadata are provided for the input sequences, the pipeline also reports the gain in taxonomic diversity in terms of the least common ancestor, together with the change in taxonomic congruence before and after amplification.

The pipeline is written in Nextflow (Di Tommaso et al., 2017), readily executable on any system with Bash and Java, publicly available at https://github.com/DessimozLab/ampliphy.

## Discussion

Throughout this study, we observed homolog enrichment consistently improves phylogenetic tree building, rather than improving alignment quality, as measured by their higher taxonomic congruence across gene families. The stronger gains we observed from the gene families from closely related taxa suggest that denser local sampling helps stabilize root placement and reduce phylogenetic error. This is consistent with prior works showing that increased taxon sampling improves phylogenetic accuracy and can mitigate systematic artifacts (Nabhan and Sarkar, 2012; Zwickl and Hillis, 2002). Importantly, these gains were achievable with a compact approach based on sequence addition, avoiding expensive full realignment. This enabled us to develop an easy-to-use pipeline that automates database search, enrichment, and downstream inference at scale.

Despite the overall trend, enrichment is not uniformly beneficial: some gene families deteriorate after adding homologs. For instance, under MAFFT L-INS-i enrichment with homologs at *E*-value *<* 10^−3^, 26 Eukaryota, 20 Amniota, 30 Bacteria, and 26 Actinomycetes HOGs out of 100 showed lower taxonomic congruence after enrichment. This can occur when highly divergent or fragmentary sequences are added, increasing alignment ambiguity (Iantorno et al., 2014; Ogden and Rosenberg, 2006). It may also occur when the addition of many taxa expands the tree search space and increases the burden of likelihood optimization (Philippe et al., 2011). In such cases, one may consider alternative methods that may complement our approach, such as species tree-aware error-correction (Wu et al., 2013).

Enrichment could also be detrimental when added sequences introduce discordance driven by biological processes such as incomplete lineage sorting (Maddison and Knowles, 2006) or horizontal gene transfer (Galtier and Daubin, 2008). In bacteria, frequent lateral gene transfer and gene turnover can produce discordance between gene history and species history (Soucy et al., 2015), which is consistent with the comparatively smaller gains we observe in bacterial datasets. Therefore, gene families that do not primarily track species evolution should be interpreted with care under enrichment.

We show that sensitive homolog sampling improves taxonomic congruence by enabling more precise root placement. However, our TCS-based rooting assessment is agnostic to cases in which rooting is difficult or potentially misleading— for example, when evolutionary rates vary strongly among branches, which can bias standard rooting strategies toward long-branch artifacts and root misplacement (Tria et al., 2017). Nevertheless, we observed the improvement is retained, even after completely removing the rooting constraints by selecting the best-scoring placement of the root. This result implies that rooting is not the sole driver of the observed gains in tree inference quality.

TCS provides a scalable way to compare topological outcomes across many gene families. We showed the validity of TCS under simulated settings, observing the identical pattern with the widely accepted Robinson-Foulds metrics. However, it is not a complete measure of phylogenetic quality, as it is topology-based, does not capture uncertainty, branch-length distortions, or event-rich histories. These might be complemented in future studies by incorporating additional axes such as branch support measures (Anisimova et al., 2011), duplication-aware (reconciliation-based) summaries (Dessimoz and Gil, 2010), and branch length–aware tree comparison criteria (Soria-Carrasco et al., 2007), among others.

TCS also inherits limitations of the reference taxonomy. For instance, we used GTDB for bacteria rather than NCBI taxonomy to improve taxonomic resolution (Supplementary Fig. 5). This implies that applying taxonomy-based evaluation is more difficult in clades with less robust or rapidly evolving classification, such as viruses (Koonin et al., 2024). Potential circularity can also arise when taxonomy is informed by genomic evidence. We mitigated this by drawing gene families and taxonomy from different resources (OMA and NCBI/GTDB, respectively), but this should be considered carefully when applying different taxonomies as a baseline.

Despite the limitations of taxonomy-based scoring, our benchmark revealed consistent trends across conditions. Our decomposition framework for evaluating alignment quality and tree inference can also serve as a practical basis for improving the underlying methods. For example, results from impoverished MSAs (Δ_*M*_ ; Fig. 2b,e), which isolate changes attributable to the aligner, allowed us to quantify that contemporary aligners outperform legacy software such as ClustalW. Likewise, using our estimated tree quality measure 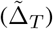, we observed higher performance of iterative maximum-likelihood tree inference compared with rapid heuristic alternatives (Supplementary Fig. 9). 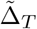 should be interpreted with caution, as it is estimated by subtracting two effects (Δ_*M*+*T*_ and Δ_*M*_ ) and can therefore be overestimated when Δ_*M*_ is underestimated (e.g., see Fig. 2b, ClustalW, *E <* 10^−10^). When interpreted with this caveat, these results suggest that our benchmarking scheme can provide useful guidance for developers pursuing more accurate and scalable phylogenomic methods.

Using this framework, we quantified the widely held expectation that dense taxon sampling benefits phylogenetic inference, and we observed that the gain arises primarily at the tree inference step. We further show that much of the improvement can be retained with a compact sequence addition strategy, enabling a scalable pipeline that automates enrichment and downstream inference. Collectively, we proposed a novel benchmarking scheme that can precisely quantify and decompose the widely known effect of denser taxon sampling, accompanied with a method that exploits this observation, providing easy-to-use solution to improve gene tree inference at scale. We expect the approach presented here to be a useful routine component of large-scale phylogenomic workflows and method assessment, particularly when additional sampling helps clarify taxonomic relationships.

## Acknowledgments

We thank Christian Ledergerber for contributing to a preliminary version of this study. We appreciate Stefano Pascarelli for constructive feedback which helped improving this work. The work was supported by Swiss National Science Foundation grants 216623 and 10005715 to C.D., and JSPS KAKENHI 16K07464 to K.K.

## Author contributions statement

D.K. and C.D. performed the analysis. D.K. developed the AmpliPhy pipeline software. M.G., K.K., and C.D. conceived the idea and conducted the preliminary study. D.K. wrote the manuscript and produced the figures, with contribution from all authors. All authors reviewed and approved the manuscript.

