## Supplementary Materials for "AmpliPhy improves gene trees by adding homologous sequences without affecting alignments"

[1]Department of Computational Biology, University of Lausanne, Lausanne, Switzerland, [2]Swiss Institute of Bioinformatics, Lausanne, Switzerland, [3]Institute of Computational Life Science, Zürich University of Applied Sciences, Wädenswil, Switzerland, [4]Department of Genome Informatics, Research Institute for Microbial Diseases, Osaka University, Suita, Japan, [5]Department of Computational Biology and Medical Sciences, Graduate School of Frontier Sciences, University of Tokyo, Kashiwa, Japan

### SUPPLEMENTARY MATERIALS

#### Figures

- Supplementary Fig. 1. Effect of homolog enrichment on alignment and tree inference in Primates gene families.
- Supplementary Fig. 2. Effect of homolog enrichment on alignment and tree inference in large Eukaryotic gene families.
- Supplementary Fig. 3. Comparison of enrichment effect across different evolutionary models.
- Supplementary Fig. 4. Effect of homolog enrichment on alignment and tree inference in bacterial gene families.
- Supplementary Fig. 5. Reduced resolution of taxonomic congruence when applying NCBI Taxonomy to bacterial species.
- Supplementary Fig. 6. Decomposed effect of enrichment by the stratification of homologs.
- Supplementary Fig. 7. Enrichment of simulated gene families improves gene tree inference towards the ground-truth.
- Supplementary Fig. 8. Taxonomic congruence score resembles Robinson-Foulds similarity from the simulated dataset.
- Supplementary Fig. 9. Iterative tree builders outperform faster heuristic alternatives.
- Supplementary Fig. 10. Tree inference remains improving after the enrichment even after removing the rooting dependency.

#### Tables

- Supplementary Table 1. Comprehensive list of OMA root HOGs used in this study.
- Supplementary Table 2. List of aligners and command lines used to run the benchmark.

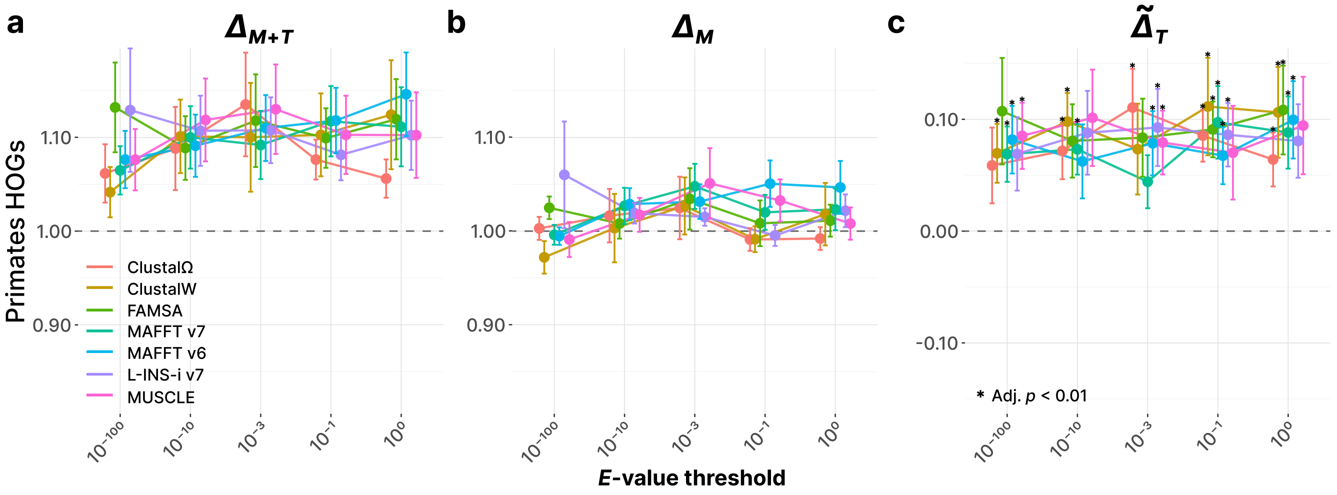


**Supplementary Fig. 1. Effect of homolog enrichment on alignment and tree inference in Primates gene families. a.** Combined effect on alignment and tree inference ($\Delta_{M+T}$) after enriching 100 HOGs from the order Primates, measured by taxonomic congruence. **b.** Effect on alignment quality ($\Delta_M$). **c.** Isolated effect on tree inference quality ($\tilde{\Delta}_T$). Statistically significant points (BH-adjusted $p < 0.01$) are annotated with asterisks.

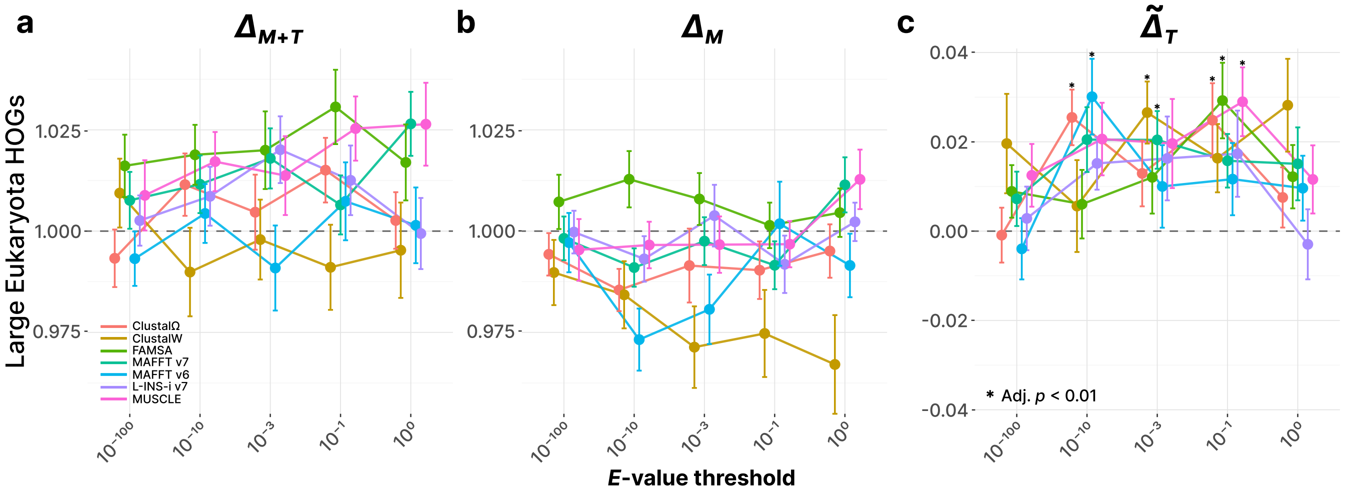


**Supplementary Fig. 2. Effect of homolog enrichment on alignment and tree inference in large Eukaryotic gene families. a.** Combined effect on alignment and tree inference ($\Delta_{M+T}$) after enriching 100 HOGs from Eukaryotes with more than 100 member sequences. **b.** Effect on alignment quality ($\Delta_M$). **c.** Isolated effect on tree inference quality ($\tilde{\Delta}_T$). Statistically significant points (BH-adjusted $p < 0.01$) are annotated with asterisks.

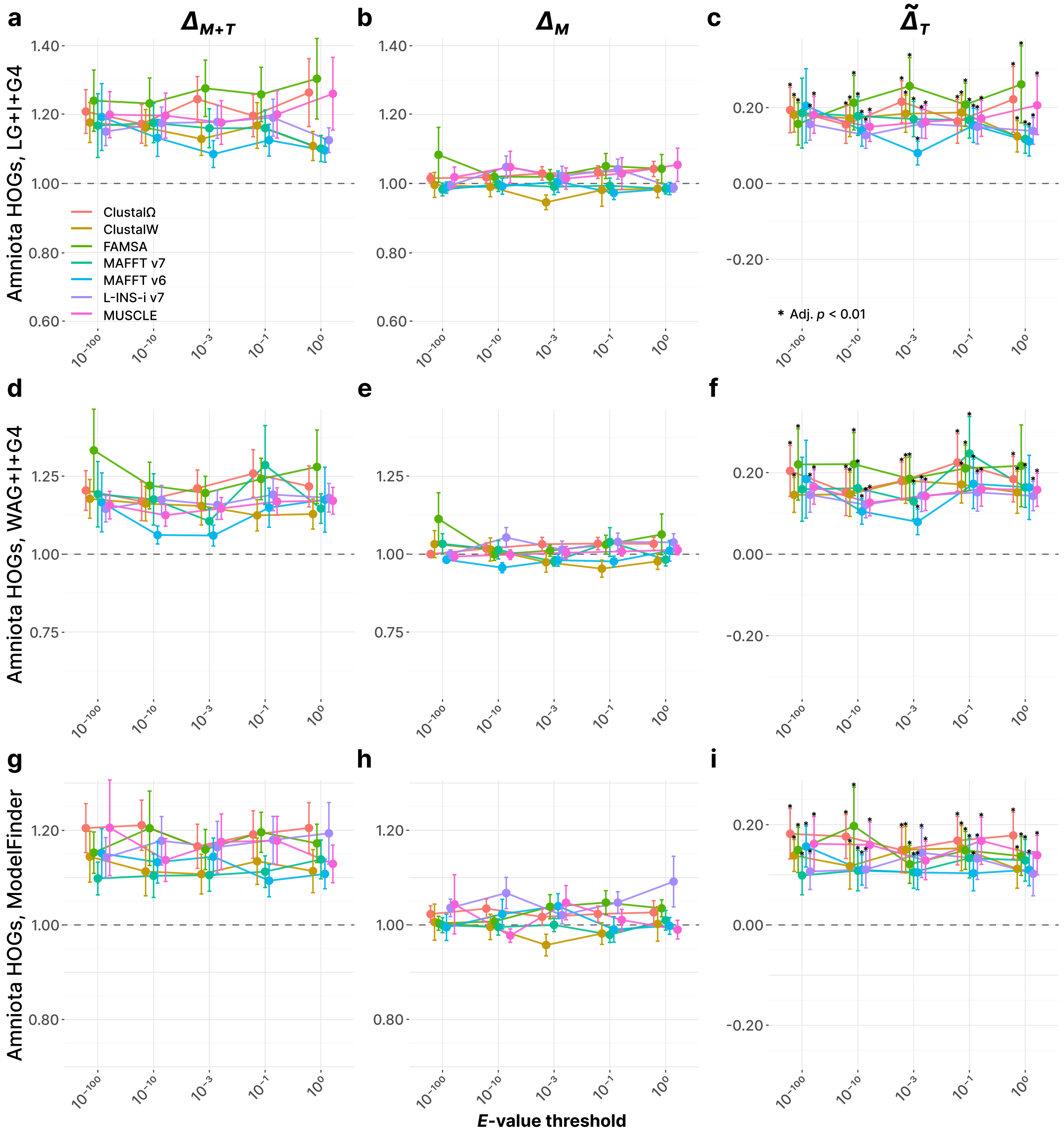


**Supplementary Fig. 3. Comparison of enrichment effect across different evolutionary models. a.** Combined effect on alignment and tree inference ($\Delta_{M+T}$) after enriching 100 HOGs from the clade Amniota. Phylogenetic trees were inferred under the model `LG+I+G4`. **b.** Effect on alignment quality ($\Delta_M$). **c.** Isolated effect on tree inference quality ($\tilde{\Delta}_T$). Statistically significant points (BH-adjusted $p < 0.01$) are annotated with asterisks. **d–f.** Corresponding data under evolutionary model `WAG+I+G4`. **g–i.** Corresponding data under the optimal model chosen by IQ-TREE 2 ModelFinder (`MF`).

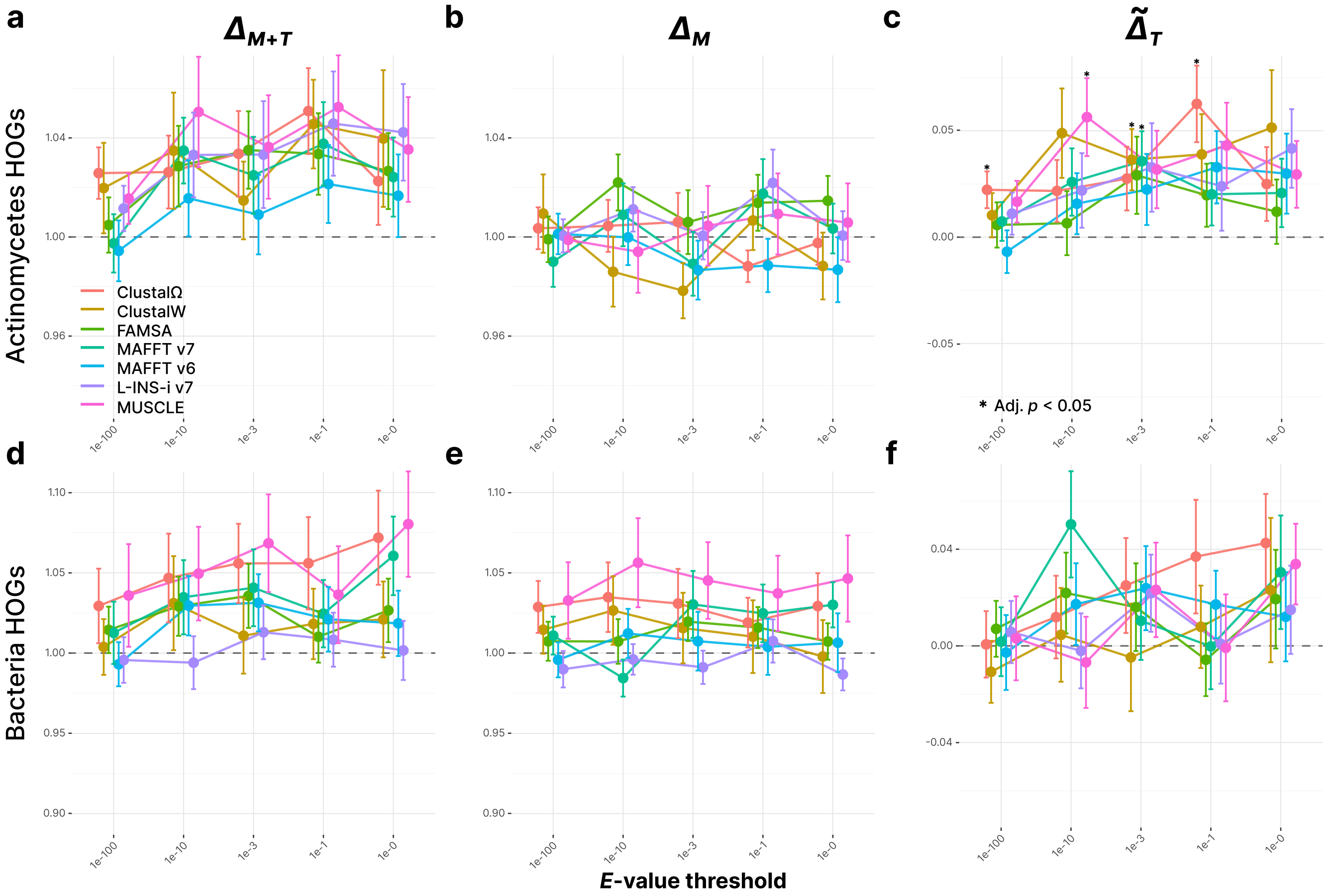


**Supplementary Fig. 4. Effect of homolog enrichment on alignment and tree inference in bacterial gene families. a.** Combined effect on alignment and tree inference ($\Delta_{M+T}$) after enriching 100 HOGs from the bacterial class Actinomycetes, measured by taxonomic congruence. **b.** Effect on alignment quality ($\Delta_M$). **c.** Isolated effect on tree inference quality ($\tilde{\Delta}_T$). Statistically significant points (BH-adjusted $p < 0.05$) are annotated with asterisks. **d–f.** Corresponding taxonomic congruence statistics for 100 HOGs spanning the entire bacterial domain.

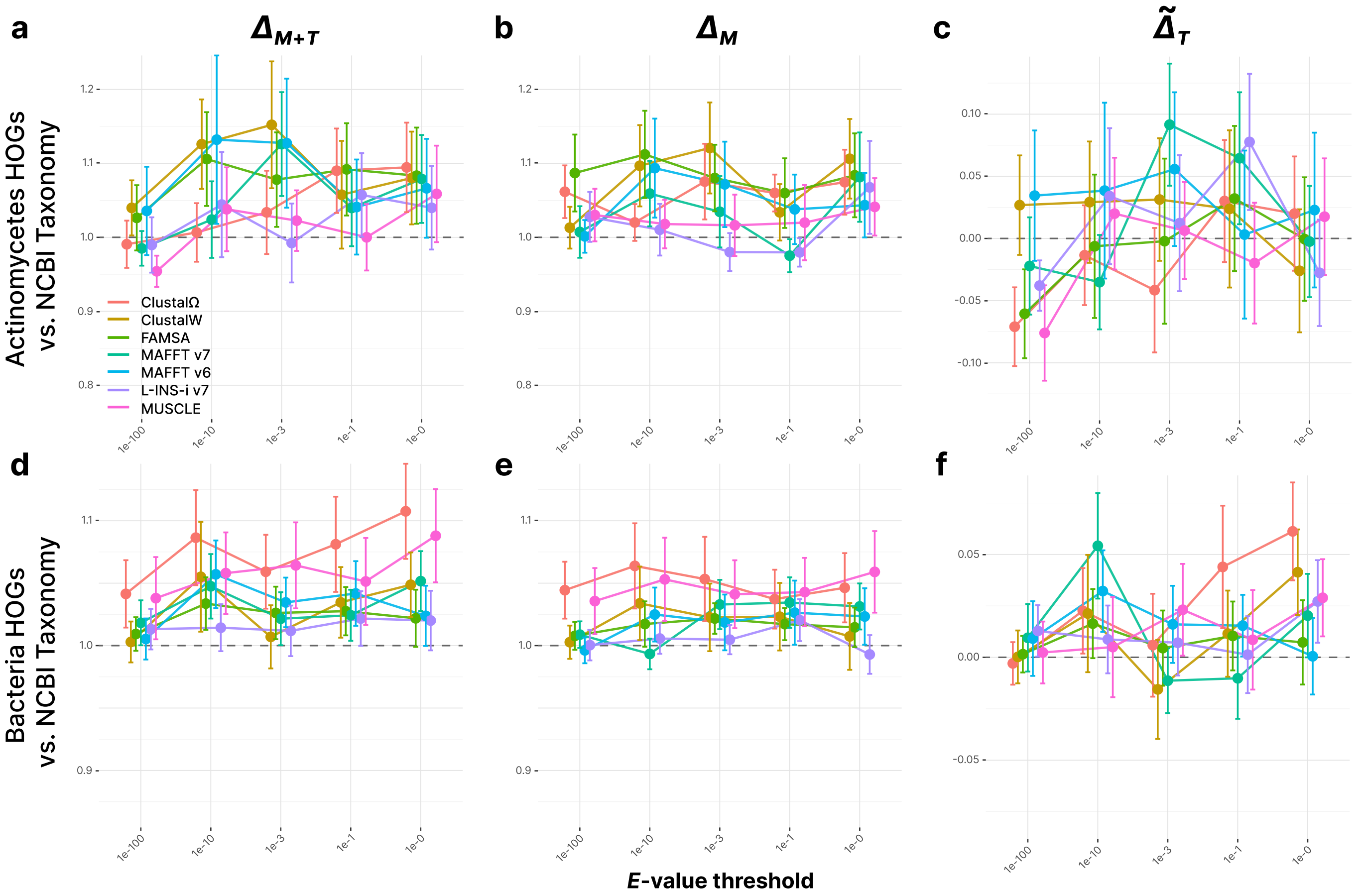


**Supplementary Fig. 5. Reduced resolution of taxonomic congruence when applying NCBI Taxonomy to bacterial species. a.** Combined effect on alignment and tree inference ($\Delta_{M+T}$) for homolog-enriched Actinomycetes HOGs, where taxonomic congruence is computed using NCBI taxonomy instead of GTDB taxonomy. **b.** Effect on alignment quality ($\Delta_M$). **c.** Isolated effect on tree inference quality ($\tilde{\Delta}_T$). **d–f.** NCBI-based taxonomic congruence statistics for 100 bacterial HOGs.

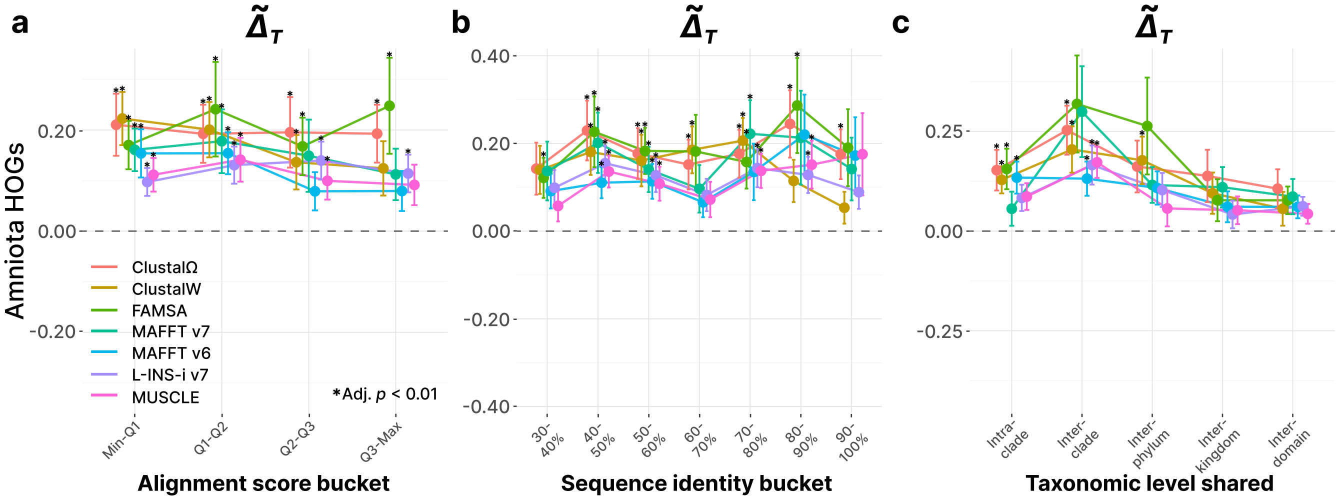


**Supplementary Fig. 6. Decomposed effect of enrichment by the stratification of homologs. a.** Impact of enrichment on the Aminota HOG tree inference quality ($\tilde{\Delta}_T$), with the homologs divided into 4 identical sized buckets based on their alignment bit scores. **b.** Impact on the tree inference with homologs grouped based on their sequence identity with respect to the query sequence ($fident$, see Methods for the definition). **c.** Impact on the tree inference with homologs grouped by their taxonomic relationship with the query clade (Amniota). Statistically significant points (BH-adjusted $p < 0.01$) are annotated with asterisks.

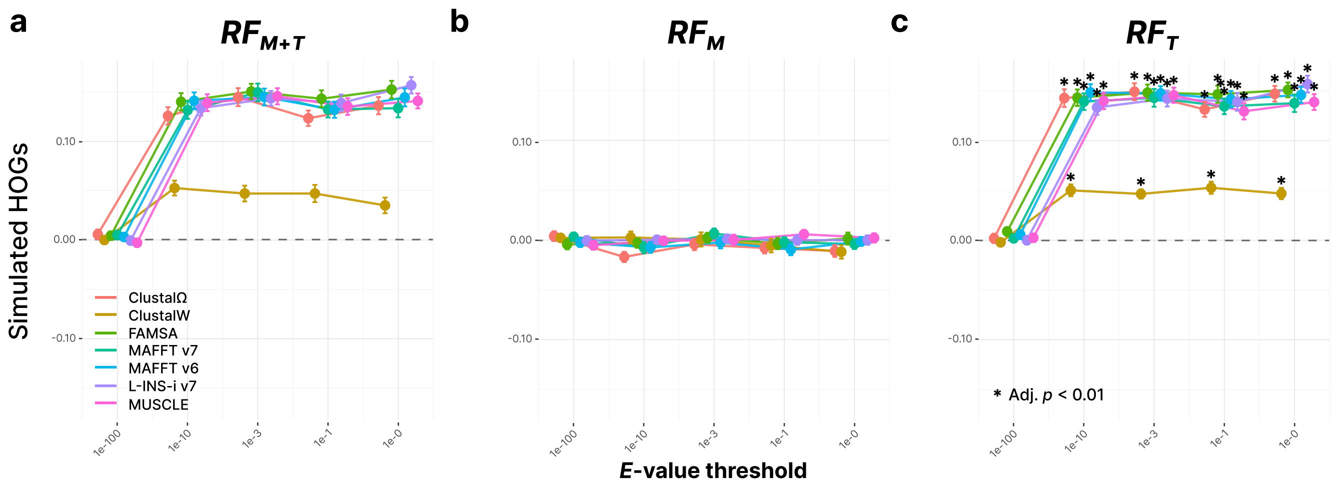


**Supplementary Fig. 7. Enrichment of simulated gene families improves gene tree inference towards the ground-truth. a.** Average Robinson-Foulds (RF) similarity (see Methods for the definition) between the ground-truth gene trees and enriched gene trees ($RF_{M+T}$), derived from the simulated evolution of 500 species with 100 gene families. The data points represent the combined impact on sequence alignment and tree inference. **b.** RF similarity between ground-truth gene trees and impoverished gene trees, representing the impact on the MSA ($RF_M$). **c.** Difference between $RF_{M+T}$ and $RF_M$, representing the isolated impact on the tree inference ($RF_T$). Statistically significant points (BH-adjusted $p < 0.01$) are annotated with asterisks.

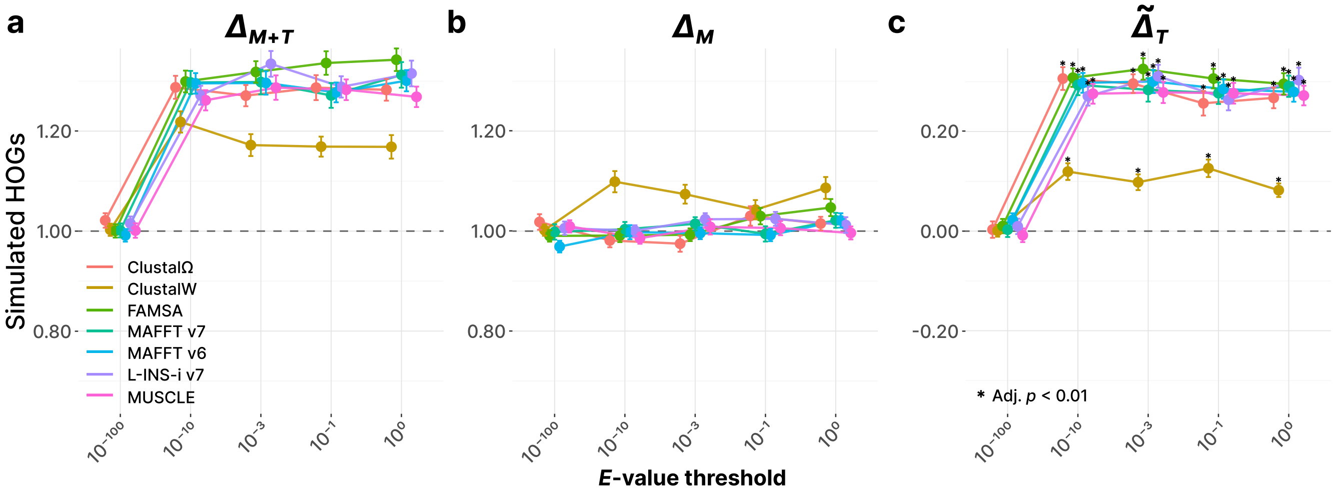


**Supplementary Fig. 8. Taxonomic congruence score resembles Robinson-Foulds similarity from the simulated dataset. a.** Combined effect on alignment and tree inference ($\Delta_{M+T}$) after enriching 100 simulated HOGs from Supplementary Fig. 7, measured by taxonomic congruence. **b.** Effect on alignment quality ($\Delta_M$). **c.** Isolated effect on tree inference quality ($\tilde{\Delta}_T$). Statistically significant points (BH-adjusted $p < 0.01$) are annotated with asterisks.

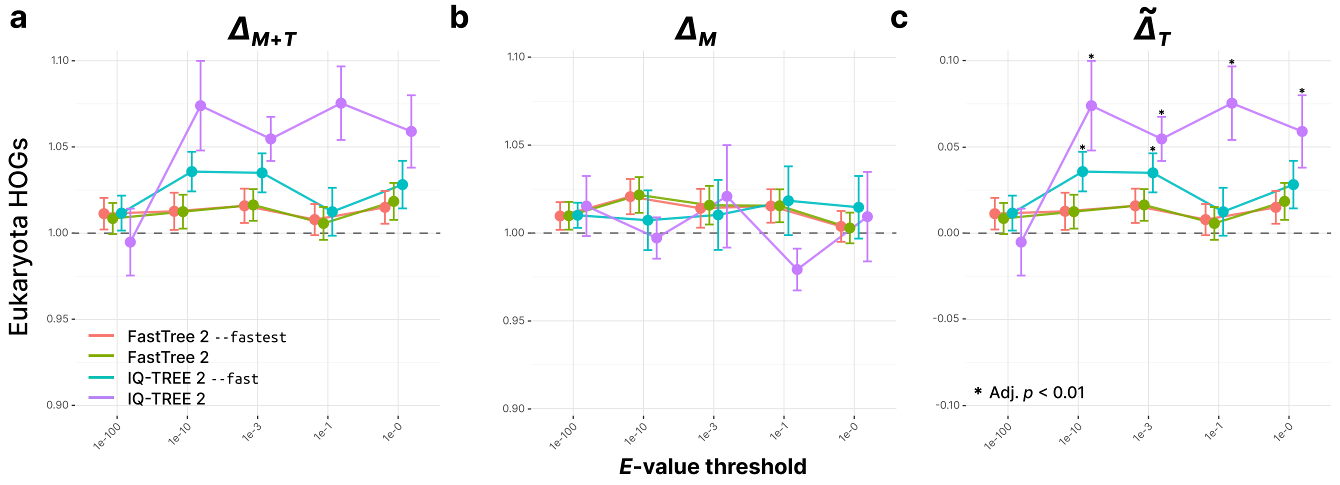


**Supplementary Fig. 9. Iterative tree builders outperform faster heuristic alternatives.** Data points are colored by the tree-building program used for phylogenetic inference, while the sequence aligner was fixed to MAFFT. **a.** Combined effect of sequence addition on alignment and tree inference ($\Delta_{M+T}$) for 100 Eukaryota HOGs. **b.** Effect on alignment quality ($\Delta_M$). **c.** Difference between $\Delta_{M+T}$ and $\Delta_M$ ($\tilde{\Delta}_T$; effect on tree inference quality). Statistically significant points (BH-adjusted $p < 0.01$) are annotated with asterisks.

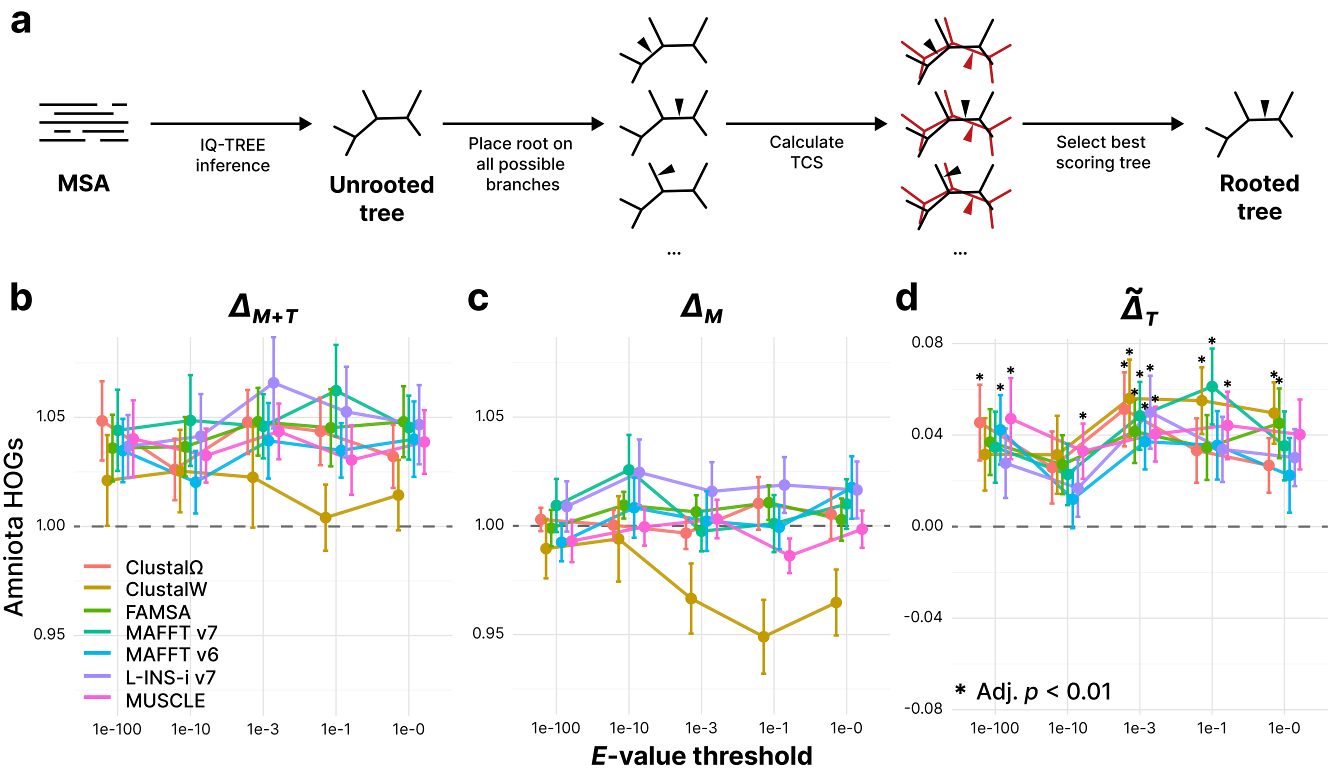


**Supplementary Fig. 10. Tree inference remains improving after the enrichment even after removing the rooting dependency. a.** Schematic illustration of the removal of rooting dependency of TCS. From the inferred unrooted tree, we calculated taxonomic congruence scores from the set of rooted trees obtained by placing root on all possible branches. Among them, we selected the rooted tree with the highest score. **b.** Combined effect on alignment and tree inference ($\Delta_{M+T}$) after enriching 100 Amniota HOGs with removed rooting dependency. **c.** Effect on alignment quality ($\Delta_M$). **d.** Isolated effect on tree inference quality ($\tilde{\Delta}_T$). Statistically significant points (BH-adjusted $p < 0.01$) are annotated with asterisks.

**Supplementary Table 1. Comprehensive list of OMA root HOGs used in this study.** Each HOG ID can be accessed in the OMA browser by querying `HOG:<HOG ID>`. The benchmark set indicates the taxon set in which each HOG was used. Taxonomic level indicates the taxon at which each HOG was inferred. Functional descriptions were retrieved from the OMA database.

| HOG ID | Benchmark set | Taxonomic level | Functional description |
|---|---|---|---|
| E0736661 | Eukaryota | Tetrapoda | Leukotriene B4 receptor |
| E0736928 | Eukaryota | Tetrapoda | KAT8 regulatory NSL complex subunit |
| E0737227 | Eukaryota | Tetrapoda | CD6 molecule |
| E0737958 | Eukaryota | Sarcopterygii | ATPase H+ transporting accessory protein |
| E0738913 | Eukaryota | Euteleostomi | NK6 homeobox |
| E0738995 | Eukaryota | Euteleostomi | BMERB domain containing |
| E0739069 | Eukaryota | Euteleostomi | HAUS augmin like complex subunit |
| E0739275 | Eukaryota | Euteleostomi | Mitogen-activated protein kinase kinase kinase 12 |
| E0739309 | Eukaryota | Euteleostomi | Myeloid-associated differentiation marker |
| E0739422 | Eukaryota | Euteleostomi | Sema domain immunoglobulin domain Ig short basic domain secreted semaphorin 3E |
| E0739485 | Eukaryota | Euteleostomi | High mobility group nucleosome-binding domain-containing protein |
| E0739504 | Eukaryota | Euteleostomi | Hydroxysteroid 17-beta dehydrogenase |
| E0739554 | Eukaryota | Euteleostomi | Angiopoietin like |
| E0739875 | Eukaryota | Euteleostomi | Macrophage scavenger receptor |
| E0741827 | Eukaryota | Euteleostomi | F-box protein 24 |
| E0745703 | Eukaryota | Gnathostomata | Matrix metalloproteinase-9 |
| E0745717 | Eukaryota | Gnathostomata | Protein-coupled receptor 20 |
| E0745736 | Eukaryota | Gnathostomata | Prostaglandin F2-alpha receptor |
| E0745773 | Eukaryota | Gnathostomata | Tumor protein p53 inducible nuclear protein |
| E0745867 | Eukaryota | Gnathostomata | Nucleotide binding oligomerization domain containing |
| E0745959 | Eukaryota | Gnathostomata | Domain containing |
| E0745969 | Eukaryota | Gnathostomata | Myozenin |
| E0746267 | Eukaryota | Gnathostomata | Forkhead box N2 |
| E0746286 | Eukaryota | Gnathostomata | EvC ciliary complex subunit |
| E0746357 | Eukaryota | Gnathostomata | Thromboxane A2 receptor |
| E0746423 | Eukaryota | Gnathostomata | Hyaluronan binding protein |
| E0746523 | Eukaryota | Gnathostomata | Family with sequence similarity 222 member |
| E0746543 | Eukaryota | Gnathostomata | Rho guanine nucleotide exchange factor |
| E0746659 | Eukaryota | Gnathostomata | Fibronectin type III domain containing |
| E0746710 | Eukaryota | Gnathostomata | Nuclear receptor interacting protein |
| E0747054 | Eukaryota | Gnathostomata | Six family member |
| E0747393 | Eukaryota | Gnathostomata | KAT8 regulatory NSL complex subunit |
| E0748047 | Eukaryota | Gnathostomata | Macrophage receptor with collagenous structure |
| E0748254 | Eukaryota | Gnathostomata | TAFA chemokine like family member |
| E0749133 | Eukaryota | Gnathostomata | DnaJ heat shock protein family Hsp40 member B2 |
| E0749136 | Eukaryota | Gnathostomata | Claudin |
| E0749142 | Eukaryota | Gnathostomata | Poly ADP-ribose polymerase |
| E0749861 | Eukaryota | Gnathostomata | Origin recognition complex subunit |
| E0752475 | Eukaryota | Vertebrata | 5-hydroxytryptamine receptor 3A |
| E0752720 | Eukaryota | Vertebrata | Fidgetin microtubule severing factor |
| E0753547 | Eukaryota | Vertebrata | Tripartite motif containing 55 |
| E0754197 | Eukaryota | Vertebrata | Lysosomal associated membrane protein |
| E0754233 | Eukaryota | Vertebrata | CACN subunit beta associated regulatory protein |
| E0756359 | Eukaryota | Chordata | ETS variant transcription factor |
| E0758308 | Eukaryota | Chordata | Gap junction protein |
| E0758684 | Eukaryota | Chordata | Collagen type IX alpha |
| E0759084 | Eukaryota | Deuterostomia | G-protein coupled receptor |
| E0759259 | Eukaryota | Deuterostomia | Peroxisome proliferator-activated receptor |
| E0759639 | Eukaryota | Deuterostomia | Family with sequence similarity 126 member |
| E0761639 | Eukaryota | Bilateria | Midnolin |
| E0769246 | Eukaryota | Bilateria | Doublecortin domain containing |
| E0770269 | Eukaryota | Bilateria | Transcription factor |
| E0778401 | Eukaryota | Eumetazoa | E3 ubiquitin-protein ligase |
| E0779894 | Eukaryota | Eumetazoa | Subcomponent-binding protein mitochondrial |
| E0780058 | Eukaryota | Eumetazoa | E2F transcription factor |
| E0780419 | Eukaryota | Eumetazoa | Partner transcriptional co-repressor |
| E0781413 | Eukaryota | Eumetazoa | Protein lin-52 homolog |
| E0781675 | Eukaryota | Eumetazoa | Derived by automated computational analysis using gene prediction method Gnomon |
| E0781703 | Eukaryota | Eumetazoa | Replication protein |
| E0781792 | Eukaryota | Eumetazoa | Protein 10 |

**Supplementary Table 1.** (Continued.)

| HOG ID | Benchmark set | Taxonomic level | Functional description |
|---|---|---|---|
| E0783173 | Eukaryota | Eumetazoa | Plexin domain containing |
| E0784066 | Eukaryota | Eumetazoa | MSL complex subunit |
| E0784079 | Eukaryota | Eumetazoa | Cell division cycle associated |
| E0785287 | Eukaryota | Eumetazoa | Trefoil factor |
| E0789185 | Eukaryota | Metazoa | Cilia and flagella associated protein 276 |
| E0789260 | Eukaryota | Metazoa | Coiled-coil domain containing 186 |
| E0790046 | Eukaryota | Metazoa | Transmembrane protein 229b |
| E0790087 | Eukaryota | Metazoa | E3 ubiquitin-protein ligase KCMF1 |
| E0790149 | Eukaryota | Metazoa | Nuclear factor of kappa light polypeptide gene enhancer in B-cells inhibitor epsilon |
| E0792964 | Eukaryota | Opisthokonta | Exosome complex component rrp45 |
| E0793021 | Eukaryota | Opisthokonta | Spindle apparatus coiled-coil protein |
| E0793757 | Eukaryota | Opisthokonta | Zinc finger hit domain-containing protein |
| E0793921 | Eukaryota | Opisthokonta | Derived by automated computational analysis using gene prediction method Gnomon |
| E0794199 | Eukaryota | Opisthokonta | Protein MIX23 |
| E0794232 | Eukaryota | Opisthokonta | Serine/threonine-protein kinase ulk3 |
| E0794379 | Eukaryota | Opisthokonta | Ankyrin repeat domain 16 |
| E0795316 | Eukaryota | Opisthokonta | Cox assembly mitochondrial protein |
| E0795420 | Eukaryota | Opisthokonta | SH2 domain containing |
| E0796129 | Eukaryota | Opisthokonta | Striatin interacting protein |
| E0801614 | Eukaryota | Eukaryota | Eukaryotic translation initiation factor 2d |
| E0801674 | Eukaryota | Eukaryota | Pescadillo homolog |
| E0801829 | Eukaryota | Eukaryota | RRNA adenine |
| E0802240 | Eukaryota | Eukaryota | Derived by automated computational analysis using gene prediction method Gnomon |
| E0802765 | Eukaryota | Eukaryota | Saccharopine dehydrogenase NADP binding domain-containing protein |
| E0803072 | Eukaryota | Eukaryota | Transmembrane protein 144 |
| E0803107 | Eukaryota | Eukaryota | Domain-containing protein |
| E0803212 | Eukaryota | Eukaryota | Thioredoxin-related transmembrane protein |
| E0803328 | Eukaryota | Eukaryota | Non-structural maintenance of chromosomes element |
| E0803575 | Eukaryota | Eukaryota | Actin related protein 2/3 complex inhibitor |
| E0803937 | Eukaryota | Eukaryota | SNARE-complex protein Syntaxin-18 N-terminal domain-containing protein |
| E0804722 | Eukaryota | Eukaryota | Coiled-coil domain containing 113 |
| E0804798 | Eukaryota | Eukaryota | DNA replication licensing factor mcm2 |
| E0804898 | Eukaryota | Eukaryota | Cilia- and flagella-associated protein 300 |
| E0805604 | Eukaryota | Eukaryota | GPI ethanolamine phosphate transferase |
| E0808137 | Eukaryota | Eukaryota | Guanine nucleotide exchange factor |
| E0812781 | Eukaryota | Eukaryota | Membrane protein |
| E0813567 | Eukaryota | Eukaryota | 6-pyruvoyl tetrahydrobiopterin synthase |
| E0816287 | Eukaryota | Eukaryota | DDB1 and CUL4 associated factor |
| E0816677 | Eukaryota | Eukaryota | Alpha subcomplex subunit 10 mitochondrial |
| E0818004 | Eukaryota | Eukaryota | TBC1 domain family member 23 |
| E0732656 | Amniota | Amniota | A-kinase anchor inhibitor |
| E0736903 | Amniota | Tetrapoda | Receptor subunit alpha |
| E0738993 | Amniota | Euteleostomi | Protein disulfide isomerase like testis expressed |
| E0739117 | Amniota | Euteleostomi | Phosphoinositide-3-kinase interacting protein |
| E0739356 | Amniota | Euteleostomi | Synapse defective Rho GTPase homolog |
| E0739363 | Amniota | Euteleostomi | Potassium channel tetramerization domain containing |
| E0740076 | Amniota | Euteleostomi | Nuclear receptor subfamily |
| E0740208 | Amniota | Euteleostomi | Myelin protein zero like |
| E0745588 | Amniota | Gnathostomata | Opsin |
| E0745662 | Amniota | Gnathostomata | Complexin |
| E0745759 | Amniota | Gnathostomata | Recombination-activating protein |
| E0746040 | Amniota | Gnathostomata | Exocyst complex component |
| E0746607 | Amniota | Gnathostomata | Dendrocyte expressed seven transmembrane protein |
| E0746839 | Amniota | Gnathostomata | Matrix metallopeptidase 23B |
| E0747039 | Amniota | Gnathostomata | C-X-C motif chemokine receptor |
| E0747049 | Amniota | Gnathostomata | Proopiomelanocortin |
| E0748381 | Amniota | Gnathostomata | Apolipoprotein A-IV |
| E0749003 | Amniota | Gnathostomata | Ataxin |
| E0749052 | Amniota | Gnathostomata | SIX homeobox |
| E0750288 | Amniota | Gnathostomata | Interleukin 17 receptor |
| E0752398 | Amniota | Vertebrata | Glucose-6-phosphatase catalytic subunit |

**Supplementary Table 1.** (Continued.)

| HOG ID | Benchmark set | Taxonomic level | Functional description |
|---|---|---|---|
| E0752660 | Amniota | Vertebrata | Zinc finger and BTB domain containing 7a |
| E0752678 | Amniota | Vertebrata | LIM and cysteine rich domains |
| E0754068 | Amniota | Vertebrata | BARX homeobox |
| E0754499 | Amniota | Vertebrata | Acetylcholine receptor subunit gamma |
| E0754546 | Amniota | Vertebrata | Paired related homeobox |
| E0755467 | Amniota | Chordata | EBF transcription factor |
| E0755569 | Amniota | Chordata | Tyrosine-protein kinase receptor |
| E0755637 | Amniota | Chordata | L-dopachrome tautomerase |
| E0756159 | Amniota | Chordata | Dopamine receptor |
| E0756270 | Amniota | Chordata | Retinol binding protein |
| E0756421 | Amniota | Chordata | Glycerophosphodiester phosphodiesterase domain containing |
| E0759640 | Amniota | Deuterostomia | ADAMTS like |
| E0760031 | Amniota | Deuterostomia | SOSS complex subunit |
| E0761363 | Amniota | Bilateria | Mediator of RNA polymerase ii transcription subunit 30 |
| E0761506 | Amniota | Bilateria | Secreted protein acidic and cysteine rich |
| E0761586 | Amniota | Bilateria | Open reading frame 120 |
| E0762353 | Amniota | Bilateria | 5-hydroxytryptamine receptor |
| E0771775 | Amniota | Bilateria | Suppressor of variegation 3-9 homolog |
| E0778191 | Amniota | Eumetazoa | ELL associated factor |
| E0778742 | Amniota | Eumetazoa | Catenin |
| E0779849 | Amniota | Eumetazoa | Serine/threonine-protein kinase 40 |
| E0779919 | Amniota | Eumetazoa | Leucine rich repeat containing 73 |
| E0780141 | Amniota | Eumetazoa | Islet cell autoantigen |
| E0780552 | Amniota | Eumetazoa | Histone acetyltransferase |
| E0781053 | Amniota | Eumetazoa | Synaptotagmin |
| E0781454 | Amniota | Eumetazoa | Mucosa-associated lymphoid tissue lymphoma translocation protein |
| E0783327 | Amniota | Eumetazoa | Zinc finger FYVE domain containing 27 |
| E0784079 | Amniota | Eumetazoa | Cell division cycle associated |
| E0789041 | Amniota | Metazoa | Transcriptional adapter |
| E0789139 | Amniota | Metazoa | Nuclear receptor binding protein |
| E0789344 | Amniota | Metazoa | Transcription initiation factor tfiid subunit |
| E0789485 | Amniota | Metazoa | Sphingosine kinase |
| E0789653 | Amniota | Metazoa | Dynactin subunit |
| E0789683 | Amniota | Metazoa | Growth arrest specific |
| E0789718 | Amniota | Metazoa | Ragulator complex protein lamtor5 |
| E0790104 | Amniota | Metazoa | RING-type E3 ubiquitin transferase |
| E0790576 | Amniota | Metazoa | Ligand of numb-protein |
| E0792967 | Amniota | Opisthokonta | Rna polymerase |
| E0792988 | Amniota | Opisthokonta | Diacylglycerol kinase |
| E0793126 | Amniota | Opisthokonta | RNA helicase |
| E0793187 | Amniota | Opisthokonta | Serine/threonine kinase |
| E0793370 | Amniota | Opisthokonta | YEATS domain-containing protein |
| E0793441 | Amniota | Opisthokonta | Transmembrane channel-like protein |
| E0793449 | Amniota | Opisthokonta | Peroxisomal biogenesis factor 11 |
| E0793477 | Amniota | Opisthokonta | Helicase lymphoid specific |
| E0793928 | Amniota | Opisthokonta | Dna damage-regulated autophagy modulator protein |
| E0794206 | Amniota | Opisthokonta | Inflammation and lipid regulator with UBA-like and NBR1-like domains |
| E0795490 | Amniota | Opisthokonta | Small ribosomal subunit protein bS6m |
| E0795765 | Amniota | Opisthokonta | Protein xrp2 |
| E0801530 | Amniota | Eukaryota | Syntaxin binding protein |
| E0801542 | Amniota | Eukaryota | Nucleoside diphosphate-linked moiety |
| E0801654 | Amniota | Eukaryota | Dolichyl-diphosphooligosaccharide–protein glycotransferase |
| E0801796 | Amniota | Eukaryota | Protein mak16 homolog |
| E0802095 | Amniota | Eukaryota | Derived by automated computational analysis using gene prediction method Gnomon |
| E0802102 | Amniota | Eukaryota | 60S ribosome subunit biogenesis protein nip7 |
| E0802214 | Amniota | Eukaryota | Reductase |
| E0802469 | Amniota | Eukaryota | Succinate dehydrogenase ubiquinone flavoprotein subunit mitochondrial |
| E0802500 | Amniota | Eukaryota | Eukaryotic translation initiation factor |
| E0802531 | Amniota | Eukaryota | Peptidyl-prolyl cis-trans isomerase |
| E0802850 | Amniota | Eukaryota | Sodium/bile acid cotransporter |
| E0803542 | Amniota | Eukaryota | Tetratricopeptide repeat |
| E0803734 | Amniota | Eukaryota | Alpha-methylacyl-coa racemase |
| E0804547 | Amniota | Eukaryota | Eukaryotic translation initiation factor 4C |

**Supplementary Table 1.** (Continued.)

| HOG ID | Benchmark set | Taxonomic level | Functional description |
|---|---|---|---|
| E0804746 | Amniota | Eukaryota | AMMECR1 domain-containing protein |
| E0804997 | Amniota | Eukaryota | Serine-threonine kinase receptor-associated protein |
| E0805071 | Amniota | Eukaryota | TRAF3-interacting protein |
| E0805082 | Amniota | Eukaryota | Acid phosphatase |
| E0805169 | Amniota | Eukaryota | WD repeat containing antisense to TP73 |
| E0805724 | Amniota | Eukaryota | ATP synthase subunit |
| E0805956 | Amniota | Eukaryota | Integrator complex subunit 14 |
| E0808060 | Amniota | Eukaryota | Dehydrogenase quinone |
| E0808147 | Amniota | Eukaryota | Mannosyl-oligosaccharide glucosidase |
| E0809946 | Amniota | Eukaryota | Leucine zipper transcription factor-like protein |
| E0810770 | Amniota | Eukaryota | F-box protein |
| E0811408 | Amniota | Eukaryota | Proteasome activator complex subunit |
| E0830763 | Amniota | Eukaryota | NADH dehydrogenase ubiquinone |
| E1027385 | Amniota | LUCA | Elongation factor |
| E1027398 | Amniota | LUCA | Glycogen starch synthase |
| E1033126 | Amniota | LUCA | Mannose-6-phosphate isomerase |
| E0722237 | Primates | Simiiformes | Zinc finger protein 528 |
| E0723174 | Primates | Primates | neuropeptide |
| E0723659 | Primates | Euarchontoglires | zinc finger protein 426 |
| E0724651 | Primates | Boreoeutheria | colipase like |
| E0724835 | Primates | Boreoeutheria | Secreted protein |
| E0727291 | Primates | Eutheria | fetal and adult testis expressed |
| E0727636 | Primates | Eutheria | zinc finger protein 529 |
| E0727690 | Primates | Eutheria | Membrane spanning 4-domains A14 |
| E0730033 | Primates | Theria | neuropeptide FF-amide peptide precursor |
| E0730175 | Primates | Theria | secreted LY6/PLAUR domain containing |
| E0730840 | Primates | Theria | natural cytotoxicity triggering receptor |
| E0731014 | Primates | Theria | Small integral membrane protein 12 |
| E0731589 | Primates | Mammalia | PABPN1 like cytoplasmic |
| E0731633 | Primates | Mammalia | family with sequence similarity 104 member |
| E0731675 | Primates | Mammalia | hemoglobin subunit mu |
| E0731702 | Primates | Mammalia | linker for activation of |
| E0731827 | Primates | Mammalia | Fc fragment of IgE receptor II |
| E0732264 | Primates | Mammalia | LLLL and CFNLAS motif containing |
| E0732280 | Primates | Mammalia | Serine rich single-pass membrane protein |
| E0732400 | Primates | Mammalia | interleukin |
| E0732604 | Primates | Amniota | spermatogenesis associated 46 |
| E0732774 | Primates | Amniota | ubiquitin specific peptidase 21 |
| E0733555 | Primates | Amniota | chromosome 19 open reading frame 38 |
| E0733700 | Primates | Amniota | zinc finger protein 12 |
| E0735286 | Primates | Amniota | interleukin-23 subunit alpha |
| E0736175 | Primates | Amniota | prickle planar cell polarity protein |
| E0736543 | Primates | Tetrapoda | natriuretic peptides |
| E0736748 | Primates | Tetrapoda | BPI fold containing family |
| E0736831 | Primates | Tetrapoda | cilia and flagella associated protein 126 |
| E0737727 | Primates | Sarcopterygii | –arginine ADP-ribosyltransferase |
| E0737776 | Primates | Sarcopterygii | transcription factor 15 |
| E0738081 | Primates | Sarcopterygii | inka box actin regulator |
| E0738139 | Primates | Sarcopterygii | BAF chromatin remodeling complex subunit BCL7C |
| E0738291 | Primates | Sarcopterygii | brain expressed associated with Nedd4 |
| E0738390 | Primates | Sarcopterygii | microfibril associated protein |
| E0739110 | Primates | Euteleostomi | zinc finger DHHC-type |
| E0739292 | Primates | Euteleostomi | S100 calcium binding protein |
| E0739299 | Primates | Euteleostomi | toll-like receptor |
| E0739722 | Primates | Euteleostomi | synapse differentiation inducing |
| E0743439 | Primates | Euteleostomi | Uncharacterized protein |
| E0745504 | Primates | Gnathostomata | single-pass membrane protein with coiled-coil domains |
| E0745593 | Primates | Gnathostomata | myozenin |
| E0745800 | Primates | Gnathostomata | phosphoinositide interacting regulator of transient receptor potential channels |
| E0745803 | Primates | Gnathostomata | phosphoinositide-3-kinase regulatory subunit |
| E0745882 | Primates | Gnathostomata | transcription factor |
| E0746007 | Primates | Gnathostomata | 5-hydroxytryptamine receptor 1A |
| E0746136 | Primates | Gnathostomata | prickle planar cell polarity protein |
| E0746397 | Primates | Gnathostomata | epidermal growth factor receptor kinase substrate 8-like protein |

**Supplementary Table 1.** (Continued.)

| HOG ID | Benchmark set | Taxonomic level | Functional description |
|---|---|---|---|
| E0746720 | Primates | Gnathostomata | family with sequence similarity 83 member |
| E0746976 | Primates | Gnathostomata | transmembrane protein 150B |
| E0746990 | Primates | Gnathostomata | profilin |
| E0747254 | Primates | Gnathostomata | LY6/PLAUR domain containing 6B |
| E0747346 | Primates | Gnathostomata | transmembrane protein 82 |
| E0747746 | Primates | Gnathostomata | proline rich |
| E0748908 | Primates | Gnathostomata | receptor protein-tyrosine kinase |
| E0748910 | Primates | Gnathostomata | cellular communication network factor |
| E0749141 | Primates | Gnathostomata | H2.0 like homeobox |
| E0749156 | Primates | Gnathostomata | complement C1s |
| E0752012 | Primates | Gnathostomata | DPY30 domain containing |
| E0752621 | Primates | Vertebrata | LY6/PLAUR domain containing |
| E0753969 | Primates | Vertebrata | ST8 alpha-N-acetyl-neuraminide alpha-2,8- sialyltransferase |
| E0754037 | Primates | Vertebrata | ferric chelate reductase |
| E0754187 | Primates | Vertebrata | serum amyloid |
| E0756044 | Primates | Chordata | solute carrier family |
| E0756517 | Primates | Chordata | coiled-coil domain containing 183 |
| E0758276 | Primates | Chordata | Kringle-containing protein marking the eye and the nose |
| E0758872 | Primates | Deuterostomia | oxidase subunit 7A2 mitochondrial |
| E0759150 | Primates | Deuterostomia | glypican |
| E0759261 | Primates | Deuterostomia | matrix AAA peptidase interacting protein |
| E0761286 | Primates | Bilateria | tubulointerstitial nephritis antigen like |
| E0761287 | Primates | Bilateria | tRNA selenocysteine 1-associated protein |
| E0761914 | Primates | Bilateria | TGF-beta activated kinase |
| E0777912 | Primates | Eumetazoa | M-phase phosphoprotein |
| E0778345 | Primates | Eumetazoa | mediator of RNA polymerase ii transcription subunit |
| E0780321 | Primates | Eumetazoa | transcription factor |
| E0781265 | Primates | Eumetazoa | spermatogenesis associated |
| E0789356 | Primates | Metazoa | factor interacting protein |
| E0789656 | Primates | Metazoa | mediator of RNA polymerase ii transcription subunit 18 |
| E0793630 | Primates | Opisthokonta | RNA polymerase III transcription initiation factor |
| E0794206 | Primates | Opisthokonta | inflammation and lipid regulator with UBA-like and NBR1-like domains |
| E0795703 | Primates | Opisthokonta | selenoprotein |
| E0797424 | Primates | Opisthokonta | TATA-box binding protein associated factor 15 |
| E0801502 | Primates | Eukaryota | isovaleryl-coa dehydrogenase mitochondrial |
| E0802195 | Primates | Eukaryota | sigma non-opioid intracellular receptor |
| E0802263 | Primates | Eukaryota | Peptidylprolyl isomerase domain and wd |
| E0802523 | Primates | Eukaryota | RNA 3'-terminal phosphate cyclase-like protein |
| E0802587 | Primates | Eukaryota | Derived by automated computational analysis using gene prediction method Gnomon |
| E0803033 | Primates | Eukaryota | ERI1 exoribonuclease |
| E0804986 | Primates | Eukaryota | protein tex261 |
| E0805984 | Primates | Eukaryota | patch domain-containing protein 11 |
| E0807183 | Primates | Eukaryota | SAP30 binding protein |
| E0808942 | Primates | Eukaryota | SCP2 domain-containing protein |
| E0811210 | Primates | Eukaryota | spermatogenesis associated |
| E0813529 | Primates | Eukaryota | N-acetyltransferase domain-containing protein |
| E0827794 | Primates | Eukaryota | Peptidase S1 domain-containing protein |
| E0827817 | Primates | Eukaryota | guanylate cyclase soluble subunit |
| E0829605 | Primates | Eukaryota | golgi-associated plant pathogenesis-related protein |
| E1027752 | Primates | LUCA | Peptidase M20 dimerisation domain-containing protein |
| E1033369 | Primates | LUCA | N-acyl-aromatic-L-amino acid amidohydrolase |
| E1033459 | Primates | LUCA | transport and golgi organization |
| E0866364 | Bacteria | Campylobacterales | Ribosomal RNA large subunit methyltransferase |
| E0866384 | Bacteria | Campylobacterales | Putative uncharacterized protein |
| E0866386 | Bacteria | Campylobacterales | Competence locus |
| E0866388 | Bacteria | Campylobacterales | Putative uncharacterized protein |
| E0866440 | Bacteria | Campylobacterales | Rare lipoprotein |
| E0866444 | Bacteria | Campylobacterales | Cell division protein |
| E0866925 | Bacteria | Campylobacteria | Putative uncharacterized protein |
| E0866981 | Bacteria | Campylobacteria | Ligase |
| E0886776 | Bacteria | Bacteroidia | Cell division protein |
| E0889623 | Bacteria | Bacteroidota | D-isomer specific 2-hydroxyacid dehydrogenase |
| E0889653 | Bacteria | Bacteroidota | Membrane protein involved in the export of O-antigen and teichoic acid |

**Supplementary Table 1.** (Continued.)

| HOG ID | Benchmark set | Taxonomic level | Functional description |
|---|---|---|---|
| E0889658 | Bacteria | Bacteroidota | Repeat protein |
| E0983296 | Bacteria | Gammaproteobacteria | Outer membrane |
| E0988205 | Bacteria | Proteobacteria | Signal peptide peptidase SppA 36K type |
| E0990658 | Bacteria | Bacteria | DNA methylase N-4/N-6 domain protein |
| E0990732 | Bacteria | Bacteria | Probable queuosine precursor transporter |
| E0990851 | Bacteria | Bacteria | Sec-independent protein translocase protein |
| E0990893 | Bacteria | Bacteria | Acetyltransferase gnat family |
| E0990905 | Bacteria | Bacteria | EscU/YscU/HrcU family type III secretion system export apparatus switch protein |
| E0990910 | Bacteria | Bacteria | Ligase |
| E0990983 | Bacteria | Bacteria | Thiamine biosynthesis protein this |
| E0991027 | Bacteria | Bacteria | Protein of unknown function DUF395 YeeE/YedE |
| E0991055 | Bacteria | Bacteria | B-type cytochrome subunit |
| E0991069 | Bacteria | Bacteria | Hydrogenase nickel insertion protein hypa |
| E0991084 | Bacteria | Bacteria | Hypothetical protein |
| E0991092 | Bacteria | Bacteria | Electron transport protein SCO1/SenC |
| E0991113 | Bacteria | Bacteria | Family protein |
| E0991463 | Bacteria | Bacteria | Outer membrane protein |
| E0991547 | Bacteria | Bacteria | C-5 cytosine-specific DNA methylase |
| E0991584 | Bacteria | Bacteria | Caax amino terminal protease family |
| E0992157 | Bacteria | Bacteria | Creatininase |
| E0992254 | Bacteria | Bacteria | Crispr-associated protein cas2 |
| E0992482 | Bacteria | Bacteria | Putative uncharacterized protein |
| E0992697 | Bacteria | Bacteria | Cobalamin biosynthesis protein |
| E0993605 | Bacteria | Bacteria | Type IV secretory pathway VirB4 |
| E0994123 | Bacteria | Bacteria | Cobalamin vitamin B12 biosynthesis CbiG protein |
| E0994427 | Bacteria | Bacteria | Alkaline phosphatase |
| E0994512 | Bacteria | Bacteria | Tap domain protein |
| E0994775 | Bacteria | Bacteria | Phytanoyl-CoA dioxygenase |
| E0995117 | Bacteria | Bacteria | Protein of unknown function DUF980 |
| E0995137 | Bacteria | Bacteria | Nitrogenase iron protein |
| E0995409 | Bacteria | Bacteria | Small basic protein |
| E0995484 | Bacteria | Bacteria | PHP domain protein |
| E0995820 | Bacteria | Bacteria | DTW domain containing protein |
| E0996090 | Bacteria | Bacteria | RNA polymerase sigma-24 subunit ECF subfamily |
| E0996091 | Bacteria | Bacteria | Taurine dioxygenase |
| E0997625 | Bacteria | Bacteria | Glycoside hydrolase family 43 |
| E0998388 | Bacteria | Bacteria | Protein of unknown function DUF839 |
| E1000672 | Bacteria | Bacteria | Protein of unknown function DUF3109 |
| E1001108 | Bacteria | Bacteria | Sec-c motif domain protein |
| E1001680 | Bacteria | Bacteria | MazG nucleotide pyrophosphohydrolase |
| E1001738 | Bacteria | Bacteria | Glycoside hydrolase family 25 |
| E1002243 | Bacteria | Bacteria | Glycoside hydrolase family 43 |
| E1008850 | Bacteria | Bacteria | Phosphate acetyltransferase |
| E1013559 | Bacteria | Bacteria | Oxidoreductase cytochrome c1 |
| E1026438 | Bacteria | Bacteria | Lyase |
| E1027257 | Bacteria | LUCA | Cell division protein ftsz |
| E1027266 | Bacteria | LUCA | Large ribosomal subunit protein uL2 |
| E1027329 | Bacteria | LUCA | Thymidine phosphorylase |
| E1027338 | Bacteria | LUCA | Methylthioribose-1-phosphate isomerase |
| E1027387 | Bacteria | LUCA | Small ribosomal subunit protein uS10 |
| E1027550 | Bacteria | LUCA | Ribosomal silencing factor RsfS |
| E1027632 | Bacteria | LUCA | Gpr1/fun34/yaah family protein |
| E1027911 | Bacteria | LUCA | Conserved hypothetical protein |
| E1027973 | Bacteria | LUCA | Dna mismatch repair protein mutl |
| E1028216 | Bacteria | LUCA | Porphobilinogen deaminase |
| E1028269 | Bacteria | LUCA | Polyphosphate kinase |
| E1028473 | Bacteria | LUCA | Queuine trna-ribosyltransferase |
| E1028477 | Bacteria | LUCA | Epoxyqueuosine reductase QueH |
| E1028560 | Bacteria | LUCA | Fructose-1,6-bisphosphatase class ii |
| E1029002 | Bacteria | LUCA | Fumarate hydratase class |
| E1029250 | Bacteria | LUCA | Potassium transport system protein kup |
| E1029441 | Bacteria | LUCA | Ethanolamine ammonia-lyase large subunit |
| E1029756 | Bacteria | LUCA | Phosphoribosyl-amp cyclohydrolase |

**Supplementary Table 1.** (Continued.)

| HOG ID | Benchmark set | Taxonomic level | Functional description |
|---|---|---|---|
| E1029773 | Bacteria | LUCA | Protein of unknown function DUF89 |
| E1030072 | Bacteria | LUCA | Redox-active disulfide protein |
| E1030201 | Bacteria | LUCA | Deoxynucleoside kinase |
| E1030314 | Bacteria | LUCA | Molybdopterin converting factor |
| E1030603 | Bacteria | LUCA | Methyltransferase |
| E1030655 | Bacteria | LUCA | Alpha alpha-trehalose-phosphate synthase |
| E1030751 | Bacteria | LUCA | Ribonuclease iii |
| E1030960 | Bacteria | LUCA | Ferrochelatase |
| E1033107 | Bacteria | LUCA | Lipid-a-disaccharide synthase |
| E1033225 | Bacteria | LUCA | 30s ribosomal protein s9 |
| E1033231 | Bacteria | LUCA | Ribosomal protein s8 |
| E1033232 | Bacteria | LUCA | 30s ribosomal protein s17 |
| E1033238 | Bacteria | LUCA | 30s ribosomal protein s12 |
| E1033252 | Bacteria | LUCA | Isocitrate dehydrogenase |
| E1033823 | Bacteria | LUCA | PSP1 domain protein |
| E1033950 | Bacteria | LUCA | Phenylalanyl-trna synthetase alpha |
| E1033955 | Bacteria | LUCA | N- 5'-phosphoribosyl anthranilate isomerase |
| E1033964 | Bacteria | LUCA | Ribosomal RNA large subunit methyltransferase |
| E1036044 | Bacteria | LUCA | Glutathionylspermidine synthase |
| E1036190 | Bacteria | LUCA | Ribosomal protein l7/l12 |
| E1036237 | Bacteria | LUCA | Formate dehydrogenase gamma subunit |
| E1036285 | Bacteria | LUCA | Protein of unknown function DUF808 |
| E1036895 | Bacteria | LUCA | Transferase |
| E1037234 | Bacteria | LUCA | DUF2238 domain-containing protein |
| E1038404 | Bacteria | LUCA | Transcription termination/antitermination protein nusA |
| E1039751 | Bacteria | LUCA | Mg chelatase subunit chli |
| E0906941 | Actinomycetes | Mycobacterium | Exported alanine and valine rich protein |
| E0908369 | Actinomycetes | Mycobacteriaceae | Conserved membrane protein of uncharacterized function |
| E0908385 | Actinomycetes | Mycobacteriaceae | Hypothetical protein |
| E0908420 | Actinomycetes | Mycobacteriaceae | Hypothetical protein |
| E0908431 | Actinomycetes | Mycobacteriaceae | Transcriptional regulator |
| E0908698 | Actinomycetes | Mycobacteriaceae | PE family protein |
| E0908753 | Actinomycetes | Mycobacteriaceae | Substrate-binding region of ABC-type glycine betaine transport system |
| E0908869 | Actinomycetes | Mycobacteriaceae | Hypothetical protein |
| E0909355 | Actinomycetes | Mycobacteriales | Transcriptional regulator LysR family |
| E0909399 | Actinomycetes | Mycobacteriales | Putative uncharacterized protein |
| E0909416 | Actinomycetes | Mycobacteriales | Two component transcriptional regulator LuxR family |
| E0909556 | Actinomycetes | Mycobacteriales | Membrane protein |
| E0909967 | Actinomycetes | Mycobacteriales | Acyltransferase |
| E0910650 | Actinomycetes | Mycobacteriales | Hypothetical protein |
| E0910862 | Actinomycetes | Actinomycetia | Putative uncharacterized protein |
| E0910923 | Actinomycetes | Actinomycetia | Putative uncharacterized protein |
| E0910958 | Actinomycetes | Actinomycetia | Major facilitator superfamily |
| E0910963 | Actinomycetes | Actinomycetia | Putative uncharacterized protein |
| E0910969 | Actinomycetes | Actinomycetia | Molybdopterin converting factor |
| E0911000 | Actinomycetes | Actinomycetia | Proton-translocating NADH-quinone oxidoreductase chain |
| E0911214 | Actinomycetes | Actinomycetia | Helix-turn-helix domain protein |
| E0911256 | Actinomycetes | Actinomycetia | Malonyl CoA-acyl carrier protein transacylase FabD2 |
| E0911917 | Actinomycetes | Actinomycetia | Putative uncharacterized protein |
| E0911951 | Actinomycetes | Actinomycetia | Membrane protein |
| E0912050 | Actinomycetes | Actinomycetia | Putative uncharacterized protein |
| E0912307 | Actinomycetes | Actinomycetia | DNA polymerase III |
| E0912826 | Actinomycetes | Actinomycetia | Roadblock/LC7 family protein |
| E0913273 | Actinomycetes | Actinomycetia | Oxidoreductase molybdopterin binding |
| E0913327 | Actinomycetes | Actinomycetia | ABC transporter |
| E0913950 | Actinomycetes | Actinomycetia | Transcriptional regulator |
| E0913985 | Actinomycetes | Actinomycetia | Transcriptional regulatory protein |
| E0914187 | Actinomycetes | Actinomycetia | Hypothetical protein |
| E0914813 | Actinomycetes | Actinomycetia | Hypothetical protein |
| E0915110 | Actinomycetes | Actinomycetia | Polyketide synthase associated protein |
| E0916679 | Actinomycetes | Actinobacteriota | ADP-ribosylation/Crystallin J1 |
| E0916784 | Actinomycetes | Actinobacteriota | Hypothetical protein |
| E0916882 | Actinomycetes | Actinobacteriota | Signal transduction histidine kinase |
| E0916891 | Actinomycetes | Actinobacteriota | PF07905 purine catabolism regulatory protein-like family |

**Supplementary Table 1.** (Continued.)

| HOG ID | Benchmark set | Taxonomic level | Functional description |
|---|---|---|---|
| E0916905 | Actinomycetes | Actinobacteriota | Membrane protein |
| E0917144 | Actinomycetes | Actinobacteriota | Putative uncharacterized protein |
| E0990835 | Actinomycetes | Bacteria | Ion transport protein |
| E0990879 | Actinomycetes | Bacteria | Rod shape-determining protein mreC |
| E0990904 | Actinomycetes | Bacteria | Ferredoxin oxidoreductase |
| E0991133 | Actinomycetes | Bacteria | Hydrogenase maturation protease |
| E0991409 | Actinomycetes | Bacteria | TRNA Ile -lysidine |
| E0991460 | Actinomycetes | Bacteria | Dihydroneopterin aldolase |
| E0992129 | Actinomycetes | Bacteria | Endonuclease |
| E0992136 | Actinomycetes | Bacteria | 5,10-methylenetetrahydrofolate reductase |
| E0992179 | Actinomycetes | Bacteria | Glutamate synthase |
| E0992182 | Actinomycetes | Bacteria | Phosphodiesterase mj0936 family |
| E0992194 | Actinomycetes | Bacteria | Putative uncharacterized protein |
| E0992444 | Actinomycetes | Bacteria | Putative uncharacterized protein |
| E0992711 | Actinomycetes | Bacteria | Hypothetical protein |
| E0992735 | Actinomycetes | Bacteria | NADH pyrophosphatase |
| E0993664 | Actinomycetes | Bacteria | N-succinyltransferase |
| E0994533 | Actinomycetes | Bacteria | MmcQ/YjbR family DNA-binding protein |
| E0995457 | Actinomycetes | Bacteria | Putative uncharacterized protein |
| E0995474 | Actinomycetes | Bacteria | Thiamin pyrophosphokinase catalytic region |
| E0996733 | Actinomycetes | Bacteria | Iron permease |
| E0998291 | Actinomycetes | Bacteria | Gas vesicle |
| E0998809 | Actinomycetes | Bacteria | Hypothetical protein |
| E1000494 | Actinomycetes | Bacteria | Putative transcriptional regulator Crp/Fnr family |
| E1001514 | Actinomycetes | Bacteria | Proteasome subunit beta |
| E1005383 | Actinomycetes | Bacteria | Nuclear transport factor |
| E1006095 | Actinomycetes | Bacteria | Hypothetical protein |
| E1007742 | Actinomycetes | Bacteria | FHA domain containing protein |
| E1008293 | Actinomycetes | Bacteria | Integration host factor |
| E1009211 | Actinomycetes | Bacteria | Putative uncharacterized protein |
| E1015623 | Actinomycetes | Bacteria | Hypothetical protein |
| E1016676 | Actinomycetes | Bacteria | Serine protease |
| E1025643 | Actinomycetes | Bacteria | N-acetylmuramoyl-L-alanine amidase |
| E1027257 | Actinomycetes | LUCA | Cell division protein ftsz |
| E1027299 | Actinomycetes | LUCA | Glycine–tRNA ligase |
| E1027489 | Actinomycetes | LUCA | 3-octaprenyl-4-hydroxybenzoate carboxy-lyase |
| E1027651 | Actinomycetes | LUCA | Phosphatidylserine decarboxylase |
| E1027665 | Actinomycetes | LUCA | Quinolinate synthetase |
| E1027669 | Actinomycetes | LUCA | L-lactate permease |
| E1027736 | Actinomycetes | LUCA | Mannose-1-phosphate guanylyltransferase |
| E1028017 | Actinomycetes | LUCA | Osmotically inducible protein |
| E1028175 | Actinomycetes | LUCA | Hydantoin racemase |
| E1028192 | Actinomycetes | LUCA | UPF0060 membrane protein |
| E1028268 | Actinomycetes | LUCA | Putative uncharacterized protein |
| E1028900 | Actinomycetes | LUCA | Glutamate racemase |
| E1029110 | Actinomycetes | LUCA | Holliday junction dna helicase ruva |
| E1029913 | Actinomycetes | LUCA | Trna delta |
| E1029981 | Actinomycetes | LUCA | Trna pseudouridine synthase |
| E1030506 | Actinomycetes | LUCA | Gcn5-related N-acetyltransferase |
| E1031052 | Actinomycetes | LUCA | Sulfate abc transporter |
| E1031550 | Actinomycetes | LUCA | Hypothetical protein |
| E1032154 | Actinomycetes | LUCA | Linocin_M18 bacteriocin protein |
| E1032317 | Actinomycetes | LUCA | Hypothetical protein |
| E1032889 | Actinomycetes | LUCA | Spoom family protein |
| E1033234 | Actinomycetes | LUCA | 50s ribosomal protein l19 |
| E1033765 | Actinomycetes | LUCA | Domain-containing protein |
| E1035591 | Actinomycetes | LUCA | Transcriptional regulator ArsR family |
| E1036199 | Actinomycetes | LUCA | Protein of unknown function DUF179 |
| E1037740 | Actinomycetes | LUCA | Hypothetical protein |
| E1039273 | Actinomycetes | LUCA | Branched-chain amino acid transport system ii carrier protein |
| E1039842 | Actinomycetes | LUCA | DNA topoisomerase |
| E1040100 | Actinomycetes | LUCA | YiaAB two helix domain protein |
| E0758622 | Eukaryota large | Chordata | cadherin |
| E0765773 | Eukaryota large | Bilateria | neurexophilin and pc-esterase domain family member |

**Supplementary Table 1.** (Continued.)

| HOG ID | Benchmark set | Taxonomic level | Functional description |
|---|---|---|---|
| E0778026 | Eukaryota large | Eumetazoa | Derived by automated computational analysis using gene prediction method Gnomon |
| E0789365 | Eukaryota large | Metazoa | glutamate receptor |
| E0792959 | Eukaryota large | Opisthokonta | t-SNARE coiled-coil homology domain-containing protein |
| E0792966 | Eukaryota large | Opisthokonta | splicing factor U2AF subunit |
| E0792972 | Eukaryota large | Opisthokonta | ectonucleoside triphosphate diphosphohydrolase |
| E0792981 | Eukaryota large | Opisthokonta | fatty acid hydroxylase domain-containing protein |
| E0792984 | Eukaryota large | Opisthokonta | u1 small nuclear ribonucleoprotein 70 kDa |
| E0793012 | Eukaryota large | Opisthokonta | ras-related protein |
| E0793013 | Eukaryota large | Opisthokonta | tyrosine-protein kinase |
| E0793020 | Eukaryota large | Opisthokonta | thioredoxin |
| E0793095 | Eukaryota large | Opisthokonta | 5'-nucleotidase domain containing |
| E0793129 | Eukaryota large | Opisthokonta | delta 3,5 -Delta 2,4 -dienoyl-CoA isomerase |
| E0793140 | Eukaryota large | Opisthokonta | sorting |
| E0793181 | Eukaryota large | Opisthokonta | transmembrane emp24 domain-containing protein |
| E0793207 | Eukaryota large | Opisthokonta | ankyrin repeat |
| E0793209 | Eukaryota large | Opisthokonta | solute carrier family 26 member |
| E0793266 | Eukaryota large | Opisthokonta | polypyrimidine tract binding protein |
| E0793469 | Eukaryota large | Opisthokonta | carboxylic ester hydrolase |
| E0793516 | Eukaryota large | Opisthokonta | major facilitator superfamily |
| E0801481 | Eukaryota large | Eukaryota | regulator of chromosome condensation |
| E0801602 | Eukaryota large | Eukaryota | Amino acid transporter transmembrane domain-containing protein |
| E0801607 | Eukaryota large | Eukaryota | Serine aminopeptidase S33 domain-containing protein |
| E0801729 | Eukaryota large | Eukaryota | charged multivesicular body protein |
| E0801760 | Eukaryota large | Eukaryota | autophagy-related protein |
| E0801764 | Eukaryota large | Eukaryota | mitochondrial-processing peptidase subunit |
| E0801815 | Eukaryota large | Eukaryota | phosphoglycerate mutase |
| E0801822 | Eukaryota large | Eukaryota | casein kinase ii subunit alpha |
| E0801833 | Eukaryota large | Eukaryota | ornithine decarboxylase |
| E0801848 | Eukaryota large | Eukaryota | nad-dependent protein |
| E0801856 | Eukaryota large | Eukaryota | U6 snRNA-associated sm-like protein lsm6 |
| E0801868 | Eukaryota large | Eukaryota | choline transporter-like protein |
| E0801870 | Eukaryota large | Eukaryota | Protein N-terminal and lysine N-methyltransferase efm7 |
| E0801887 | Eukaryota large | Eukaryota | replication factor |
| E0801902 | Eukaryota large | Eukaryota | Cyclin |
| E0801907 | Eukaryota large | Eukaryota | Large ribosomal subunit protein uL23 |
| E0801918 | Eukaryota large | Eukaryota | Derived by automated computational analysis using gene prediction method Gnomon |
| E0801941 | Eukaryota large | Eukaryota | RBR-type E3 ubiquitin transferase |
| E0801945 | Eukaryota large | Eukaryota | NADH-cytochrome b5 reductase |
| E0801960 | Eukaryota large | Eukaryota | cornichon family AMPA receptor auxiliary protein |
| E0801973 | Eukaryota large | Eukaryota | peptidylprolyl isomerase |
| E0802012 | Eukaryota large | Eukaryota | serine palmitoyltransferase long chain base subunit |
| E0802046 | Eukaryota large | Eukaryota | ethanolamine kinase |
| E0802055 | Eukaryota large | Eukaryota | Derived by automated computational analysis using gene prediction method Gnomon |
| E0802134 | Eukaryota large | Eukaryota | AAA+ ATPase domain-containing protein |
| E0802137 | Eukaryota large | Eukaryota | rna helicase |
| E0802203 | Eukaryota large | Eukaryota | cystathionine |
| E0802221 | Eukaryota large | Eukaryota | cell cycle control protein |
| E0802233 | Eukaryota large | Eukaryota | serine incorporator |
| E0802298 | Eukaryota large | Eukaryota | charged multivesicular body protein |
| E0802319 | Eukaryota large | Eukaryota | defective in cullin neddylation |
| E0802323 | Eukaryota large | Eukaryota | Magnesium transporter |
| E0802328 | Eukaryota large | Eukaryota | serine/threonine-protein phosphatase 2a 55 kda regulatory subunit |
| E0802335 | Eukaryota large | Eukaryota | equilibrative nucleoside transporter |
| E0802337 | Eukaryota large | Eukaryota | Ubiquitin-like domain-containing protein |
| E0802395 | Eukaryota large | Eukaryota | casein kinase ii subunit beta |
| E0802409 | Eukaryota large | Eukaryota | Sodium/calcium exchanger membrane region domain-containing protein |
| E0802418 | Eukaryota large | Eukaryota | vesicle-associated membrane protein |
| E0802421 | Eukaryota large | Eukaryota | V-type proton ATPase subunit |
| E0802457 | Eukaryota large | Eukaryota | peroxisomal membrane protein |
| E0802506 | Eukaryota large | Eukaryota | 60s ribosomal protein l21 |
| E0802572 | Eukaryota large | Eukaryota | ribonucleoside-diphosphate reductase subunit m2 |

**Supplementary Table 1.** (Continued.)

| HOG ID | Benchmark set | Taxonomic level | Functional description |
|---|---|---|---|
| E0802609 | Eukaryota large | Eukaryota | pyruvate decarboxylase |
| E0802642 | Eukaryota large | Eukaryota | t-SNARE coiled-coil homology domain-containing protein |
| E0802656 | Eukaryota large | Eukaryota | carboxylic ester hydrolase |
| E0802701 | Eukaryota large | Eukaryota | oxidase |
| E0802750 | Eukaryota large | Eukaryota | Derived by automated computational analysis using gene prediction method Gnomon |
| E0802759 | Eukaryota large | Eukaryota | TatD DNase domain containing |
| E0802823 | Eukaryota large | Eukaryota | ABC1 atypical kinase-like domain-containing protein |
| E0802949 | Eukaryota large | Eukaryota | amine oxidase |
| E0802977 | Eukaryota large | Eukaryota | Derived by automated computational analysis using gene prediction method Gnomon |
| E0803038 | Eukaryota large | Eukaryota | cysteine and glycine-rich protein |
| E0803087 | Eukaryota large | Eukaryota | Amidase domain-containing protein |
| E0803398 | Eukaryota large | Eukaryota | CSC1/OSCA1-like 7TM region domain-containing protein |
| E0803403 | Eukaryota large | Eukaryota | Derived by automated computational analysis using gene prediction method Gnomon |
| E0803479 | Eukaryota large | Eukaryota | RING-type E3 ubiquitin transferase |
| E0804281 | Eukaryota large | Eukaryota | protein-serine/threonine phosphatase |
| E0804434 | Eukaryota large | Eukaryota | Serine/threonine protein phosphatase 2a regulatory subunit |
| E0804483 | Eukaryota large | Eukaryota | C2H2-type domain-containing protein |
| E0804978 | Eukaryota large | Eukaryota | protein yippee-like |
| E0805099 | Eukaryota large | Eukaryota | Bidirectional sugar transporter sweet |
| E0805279 | Eukaryota large | Eukaryota | phosphotransferase |
| E0805484 | Eukaryota large | Eukaryota | Derived by automated computational analysis using gene prediction method Gnomon |
| E0805752 | Eukaryota large | Eukaryota | domain-containing protein |
| E0805789 | Eukaryota large | Eukaryota | chloride intracellular channel protein |
| E0805822 | Eukaryota large | Eukaryota | voltage-dependent anion-selective channel protein |
| E0806548 | Eukaryota large | Eukaryota | domain-containing protein |
| E0809154 | Eukaryota large | Eukaryota | UDP-glucuronosyltransferase |
| E0810427 | Eukaryota large | Eukaryota | chitinase |
| E0811656 | Eukaryota large | Eukaryota | annexin |
| E1027260 | Eukaryota large | LUCA | ribose-phosphate pyrophosphokinase |
| E1027275 | Eukaryota large | LUCA | Small ribosomal subunit protein uS5 |
| E1027281 | Eukaryota large | LUCA | proteasome subunit alpha type |
| E1027363 | Eukaryota large | LUCA | Developmentally regulated gtp binding protein |
| E1027592 | Eukaryota large | LUCA | L-lactate dehydrogenase |
| E1027833 | Eukaryota large | LUCA | Enoyl reductase ER domain-containing protein |
| E1028318 | Eukaryota large | LUCA | solute carrier family 23 member |
| E1028583 | Eukaryota large | LUCA | FMN hydroxy acid dehydrogenase domain-containing protein |
| E1028629 | Eukaryota large | LUCA | ammonium transporter rh type |

**Supplementary Table 2. List of sequence aligners and command lines used to run the benchmark.** Each aligner name corresponds to the legend in **Fig. 2a**. Options used to set the number of CPU threads are omitted.

| Aligner name | Version | Command line |
|---|---|---|
| ClustalΩ | v1.2.4 | `clustalo -i <input.fa>` |
| ClustalW | v2.1 | `clustalw2 -INFILE=<input.fa> -QUIET -OUTPUT=FASTA -TYPE=PROTEIN -OUTFILE=<output.fa> &>/dev/null` |
| FAMSA | v2.2.3 | `famsa <input.fa> STDOUT 2>/dev/null` |
| MAFFT v7 | v7.526 | `mafft --anysymbol --quiet <input.fa>` |
| MAFFT v6 | v6.864 | `mafft --anysymbol --quiet <input.fa>` |
| L-INS-i v7 | v7.526 | `mafft-linsi --anysymbol --quiet <input.fa>` |
| MUSCLE | v5.3 | `muscle -align <input.fa> -output /dev/stdout 2>/dev/null` |
